# Investigating tissue-relevant causal molecular mechanisms of complex traits using probabilistic TWAS analysis

**DOI:** 10.1101/808295

**Authors:** Yuhua Zhang, Corbin Quick, Ketian Yu, Alvaro Barbeira, The GTEx Consortium, Francesca Luca, Roger Pique-Regi, Hae Kyung Im, Xiaoquan Wen

## Abstract

Transcriptome-wide association studies (TWAS), an integrative framework using expression quantitative trait loci (eQTLs) to construct proxies for gene expression, have emerged as a promising method to investigate the biological mechanisms underlying associations between genotypes and complex traits. However, challenges remain in interpreting TWAS results, especially regarding their causality implications. In this paper, we describe a new computational framework, probabilistic TWAS (PTWAS), to detect associations and investigate causal relationships between gene expression and complex traits. We use established concepts and principles from instrumental variables (IV) analysis to delineate and address the unique challenges that arise in TWAS. PTWAS utilizes probabilistic eQTL annotations derived from multi-variant Bayesian fine-mapping analysis conferring higher power to detect TWAS associations than existing methods. Additionally, PTWAS provides novel functionalities to evaluate the causal assumptions and estimate tissue- or cell-type specific causal effects of gene expression on complex traits. These features make PTWAS uniquely suited for in-depth investigations of the biological mechanisms that contribute to complex trait variation. Using eQTL data across 49 tissues from GTEx v8, we apply PTWAS to analyze 114 complex traits using GWAS summary statistics from several large-scale projects, including the UK Biobank. Our analysis reveals an abundance of genes with strong evidence of eQTL-mediated causal effects on complex traits and highlights the heterogeneity and tissue-relevance of these effects across complex traits. We distribute software and eQTL annotations to enable users performing rigorous TWAS analysis by leveraging the full potentials of the latest GTEx multi-tissue eQTL data.

## 1 Introduction

Over the past two decades, genome-wide association studies (GWAS) have identified an abundance of genetic loci associated with complex traits and diseases [1]. However, most GWAS loci are located in the non-coding regions of the genome, and our understanding of the mechanisms and causal relationships underlying these associations is still lacking. Direct experimental investigation of the causal path from genotype to complex trait (e.g., via randomized controlled experiments on human subjects using CRISPR) is limited by technical difficulties and ethical constraints. Thus, statistical methods for causal inference from observational data become a logical alternative, especically with the increasing availability of large-scale GWAS and molecular QTL data. Instrumental variable (IV) analysis is an established inference framework to investigate causal relationships from observational data in the presence of possible confounding, which has been widely applied in econometrics and epidemiology. In observational epidemiology, the study of health consequences of modifiable environmental factors by utilizing genetic variants as instrumental variables is commonly known as Mendelian randomization (MR), which has shown great promise when active intervention is implausible [2, 3].

IV analysis naturally integrates genetic, molecular phenotype, and complex trait data and investigates the causal links from molecular phenotypes (e.g., gene expressions, DNA methylations) leading to complex diseases. Most popular approaches for transcriptome-wide association analysis (TWAS), e.g., PrediXcan [4], TWAS-Fusion [5], and SMR [6], all can be viewed as some forms of IV analysis with an emphasis on *testing* causal relationships from gene expressions to complex traits. IV analysis, in comparison to other competing causal inference frameworks, is particularly suitable for TWAS because of its explicit consideration of confounding factors in the model formulation. However, we note that existing TWAS methods have at least two notable weaknesses when considering under the framework of IV analysis. First, validating the causal implications from the positive association findings from TWAS is mostly unaddressed. Second, efficient *estimation* of causal effects remains an open problem. The latter issue is particularly crucial for investigating causal molecular mechanisms in different cellular environments. In this paper, we show that the solutions to these problems can be directly derived from the established principles of IV analysis.

Compared to the traditional applications of IV/MR analysis, TWAS also presents some unique challenges. Most notably, it deals with tens of thousands of candidate genes for each given complex trait of interest, which warrants a powerful and efficient testing procedure. Such procedures are under-developed in the traditional IV analysis literature. Additionally, it is now well-known that a single gene can be regulated by multiple independent genetic variants, which is known as allelic heterogeneity (AH) [7]. However, the available molecular QTLs for any given gene are quite limited compared to the conventional MR analysis. This feature prevents direct applications of some established analytic approaches like Egger regression [8], which requires a relatively large number of independent eQTLs. Finally, both molecular QTLs and GWAS hits are inferred from association analysis and embedded with uncertainty at single variant levels due to linkage disequilibrium (LD). Simultaneous accounting for allelic heterogeneity and LD in TWAS will likely significantly improve the precision and power of the analysis. This issue also remains unsolved in a computationally efficient way.

In this paper, we propose a new analytic framework named as *probabilistic analysis of transcriptome-wide association study*, or PTWAS, to address three interrelated areas in TWAS analysis:

1. Testing of the causal relationship from genes to complex traits
2. Validating causal implications through model diagnosis
3. Estimating causal effects from genes to complex traits (i.e., gene-to-trait effects)

PTWAS is built upon the causal inference framework of IV analysis and is designed specifically to deal with the challenges arising from the TWAS analysis. PTWAS utilizes the probabilistic annotations of eQTLs derived from the state-of-the-art multi-SNP analysis method for fine-mapping of *cis*-eQTLs. This feature provides excellent convenience and flexibility to account for uncertainty in eQTLs for both testing and estimation purposes. Taking advantage of wide-spread allelic heterogeneity, PTWAS also provides a principled way to examine the causal implications from the PTWAS analysis. Through theoretical derivation and numerical simulation studies, we illustrate the advantages of PTWAS over the existing methods. We apply PTWAS to analyze the eQTL data from the GTEx project (version 8) [9] and 114 complex traits from the UK Biobank [10] and other large-scale consortia. Our results demonstrate the unique benefits of utilizing estimated gene-to-trait effects (rather than testing *p*-values) in investigating and interpreting causal molecular mechanisms of complex traits. The software package and utilities implementing the proposed computational approaches and relevant resources are available in https://github.com/xqwen/ptwas.

## 2 Results

### 2.1 Method overview

The causal inference procedure implemented in PTWAS involves three sequential stages that cover hypothesis testing, causal assumption validation, and effect size estimation. PTWAS works with summary-level statistics from GWAS and probabilistic annotations of eQTLs derived from the multi-SNP fine-mapping analysis. The PTWAS procedure starts with a multiple hypothesis testing of causal relations between candidate genes and the complex trait of interest, which yields an outcome that is similar to the results of PrediXcan and TWAS-Fusion. For all the genes rejected in the first stage, PTWAS further performs a model diagnosis procedure to detect potential violations of the critical causal assumptions in the IV analysis. Finally, an estimate of the causal effect on the complex trait of interest is derived for each qualified gene that passes previous screening steps. Figure 1 summarizes the workflow of the complete PTWAS procedure.

**Figure 1:**
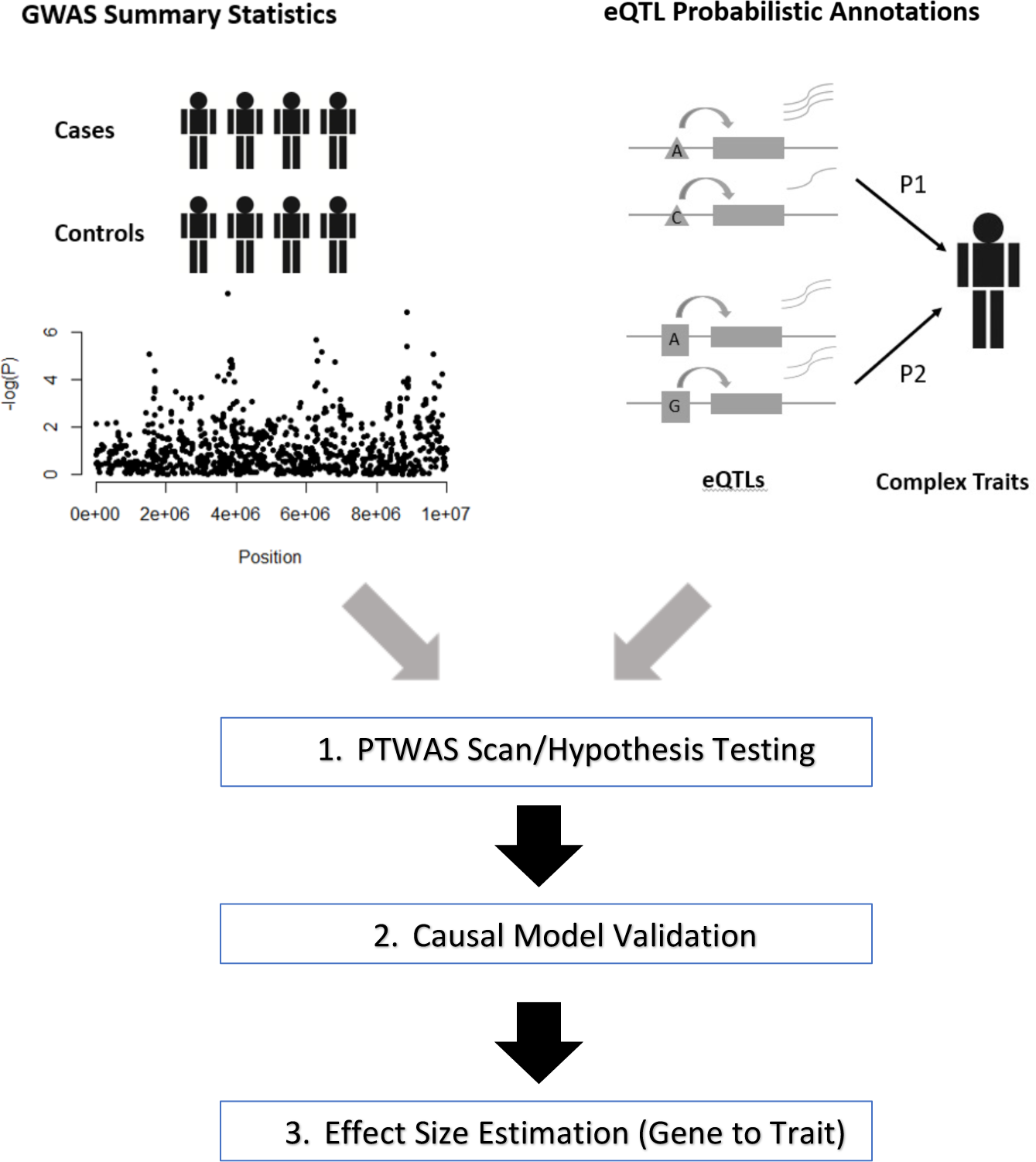
Workflow of PTWAS analysis.

A key feature of PTWAS is that we adopt a well-established statistical principle in the IV analysis to construct and utilize *strong* instruments for inference and model diagnosis. Specifically, we use the probabilistic quantifications of *cis*-eQTLs derived from the fine-mapping analysis to characterize the strength of eQTLs while accounting for allelic heterogeneity and LD [11, 12]. For each gene, the adopted probabilistic annotations provide comprehensive information at three different levels: i) the posterior probability for each plausible association model (i.e., the combination of genetic variants), which we refer to as the *posterior model probability*; ii) the posterior inclusion probability (PIP) for each SNP (or, SNP-level PIP); and iii) the posterior probability for an independent signal cluster, which we refer to as signal-level PIP, or SPIP. A signal cluster consists of a set of SNPs in LD that are deemed to represent the same underlying causal eQTL (but not completely identifiable based on the association data). The Bayesian credible set now commonly used in fine-mapping analyses of genetic associations [13] is special cases of signal clusters. (i.e., a (100 × *p*)% Bayesian credible set can be constructed from a signal cluster by selecting the minimum subset of representing SNPs with cumulative PIPs ≥ *p*.)

#### Multiple hypothesis testing

For hypothesis testing, we construct a *composite instrumental variable* [14, 15] as the test statistic by combining the strength of multiple independent eQTLs from a given gene. Specifically, we first derive a standard linear least-square prediction function of expression phenotype for each plausible genetic association model identified from the eQTL fine-mapping analysis. We then compute the composite IV as the weighted sum of the resulting least-square predictions, where the individual weight is computed from the corresponding posterior model probability. The composite IV remains a linear function of genotypes and is subsequently used to test correlation with the complex trait of interest. Henceforward, we refer to the multiple testing procedure between constructed composite IVs and the complex trait as the PTWAS scan. Same to the available TWAS methods like PrediXcan and TWAS-Fusion, the proposed hypothesis testing procedure aims to qualitatively examine the presence/absence of the causal relationships from candidate genes to the complex trait of interest. The difference between the methods lies in the way that the composite IV is constructed. It is crucial to note that the implication from these scan procedures is limited: it provides interpretations on neither the direction nor the magnitude of the underlying gene-to-trait effects if it asserts causal relationships exist [16].

#### Validating causal assumption

For the genes that pass the initial PTWAS scan, we consider validating the causal assumptions of the IV analysis and further estimating the causal effects from implicated genes to the complex trait. The intuition for the proposed procedure is that each independent eQTL should provide an independent estimate of the *same* gene-to-trait effect under the assumptions of IV analysis. Consequently, the estimates of the effect from multiple independent eQTLs are expected to be highly consistent if the causal assumption of exclusion restriction (ER) holds. Conversely, a significant degree of inconsistency/heterogeneity observed from the estimates indicates potentially severe violations of the causal assumption. To implement this idea, we first construct an eQTL-level estimate of the gene-to-trait effect for each *strong* eQTL signal cluster (by thresholding on the signal-level PIPs). The emphasis on exclusive usage of strong eQTLs is because weak eQTLs are considered to be weak instruments, which provides unreliable and severely biased causal effect estimates. Within each selected signal cluster, we first compute a SNP-level estimate, known as the Ward ratio estimate, for each member SNP. We then combine the SNP-level estimates from the member SNPs weighted by their corresponding PIPs. This novel signal cluster-level estimator is derived from the principle of Bayesian model averaging (BMA), and fully accounts for the uncertainty by LD. Across multiple strong eQTL signals, we compute an *I*^2^ statistic [17] to quantify the heterogeneity of those independent estimates. The possible values of an *I*^2^ statistic range from 0 to 1, with *I*^2^ = 1, indicating the highest level of inconsistency.

#### Causal effect estimation

For the genes with small *I*^2^ value, we further aggregate the signal-level estimates by applying a fixed-effect meta-analysis model, which results in a final gene-level estimate of the causal effect by PTWAS. This estimator of gene-to-trait effects is similar to the IVW estimator in the IV analysis literature [15], which is known to be robust and efficient [18]. However, there is an important distinction: our aggregation of the estimates of individual instruments occurs at the levels of independent eQTL clusters rather than individual SNPs. Furthermore, we only admit strong eQTLs (judging by their signal-level PIPs) to the estimation procedure. These new features likely make the proposed estimator even more efficient.

PTWAS conceptually connects to all the existing TWAS methods, especially in the stage of scanning candidate genes. However, it also presents some notable distinctions. Both PTWAS and SMR are derived from the IV analysis. However, PTWAS explicitly takes advantage of the wide-spread allelic heterogeneity by constructing the composite IV and testing causal effects from multiple independent eQTLs. Both TWAS-Fusion and PrediXcan focus on *predicting/imputing* gene expression levels from eQTL information. The composite IVs constructed by PTWAS can also be interpreted as a natural *ensemble* predictions of gene expressions. Additionally, the separation of the hypothesis testing and the effect size estimation procedures is also a unique feature in PTWAS. Particularly, the emphasis on estimation accuracy prompts a slightly different statistical strategy from hypothesis testing. Our proposed estimation and model validation procedures are derived from the well-established principles of the IV analysis and should provide unbiased estimates and calibrated error quantifications while taking advantage of widespread allelic heterogeneity and accounting for LD. Most importantly, these properties enable valid and meaningful comparisons of gene-to-trait effects in different cellular environments within the context of TWAS.

### 2.2 Improved power for testing causal genes

We perform simulation studies to validate the proposed PTWAS method and compare its performance to the existing approaches. Our simulated data sets attempt to match the features of observed eQTL and GWAS data in practice, e.g., the distribution of *z*-statistics from single-SNP analysis, by including both strong sparse association signals and weak polygenic background effects.

We compare the PTWAS scan procedure to the existing methods, including PrediXcan, TWAS-Fusion, and SMR. We find that all examined approaches effectively control the type I errors in the scan procedure of testing causal genes (Figure 2). However, approaches explicitly accounting for allelic heterogeneity and utilizing multiple independent eQTLs (i.e., PTWAS, TWAS-Fusion, and PrediXcan) show much-improved power over SMR (which utilizes only a single instrument). Among the three multi-SNP TWAS methods, PTAWS consistently exhibits the highest power under our simulation settings, which is shown by the ROC curves in Figure 2.

**Figure 2:**
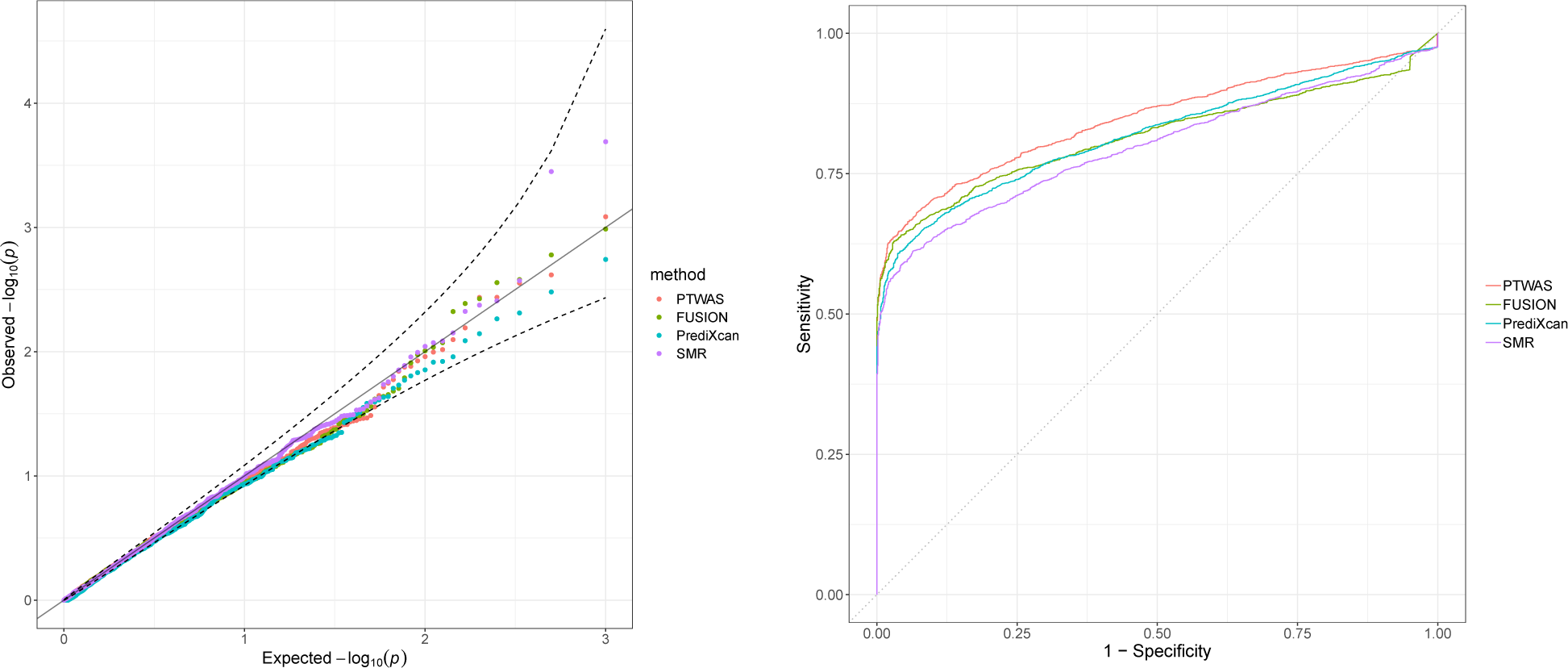
Type I error control and power in hypothesis testing of causal relationships by various methods in TWAS. The left panel shows the QQ-plot of testing *p*-values as the simulated data are specifically generated from the null scenario. All methods properly control the type I errors. The right panel shows the receiver operating characteristic (ROC) curves by different methods when the simulated data are from a mixture of null and alternative scenarios. At any specified value of the false-positive rate, PTWAS shows the highest true-positive rate among compared methods. All methods utilizing multiple independent eQTLs show higher powers than SMR (which uses a single best eQTL of each gene as the corresponding instrument).

Using the simulated data, we inspect the weight distribution over SNPs from the composite IVs constructed by different approaches. Specifically, we measure the concentration/dispersion of the (absolute) weights among SNPs using the Gini coefficient, which effectively measures the sparsity of weights [19]). Particularly, a Gini coefficient → 1 indicates a highly sparse weight distribution, i.e., only a few SNPs play essential roles in the corresponding composite IVs; whereas a Gini coefficient → 0 indicates the weights are evenly distributed in all candidate SNPs. The distributions of Gini coefficients computed from simulated datasets by PTWAS, PrediXcan, and PTWAS are shown in Supplementary Figure S2. We find that weights constructed by TWAS-Fusion are most dispersed and the weights by PrediXcan and PTWAS are mostly highly concentrated (with the mean of the Gini coefficient across datasets nearing 1). Compared to PrediXcan, the distribution of the Gini coefficients from PTWAS is mostly concentrated but shows a notably extended left tail representing dispersed weight allocation. Upon further inspection, we find those cases correspond to the scenarios where the causal SNPs lie in high LD regions. From the fundamental principle of the IV analysis, only strong instruments should be considered, which suggests an overall sparse weighting scheme. On the other hand, SNPs in high LD are not distinguishable and should receive similar weights. Among the methods tested, only the weighting scheme from PTWAS satisfies both desired properties.

The observations from this set of simulations seem to confirm that our design principle of PTWAS for explicitly accounting for allelic heterogeneity and LD contributes to the improved scan performance.

### 2.3 Accurate estimation of causal effects

We further examine the estimation procedure by PTWAS and various TWAS methods using simulations. In this particular experiment, we focus on estimating the effects of genes that pass the initial PTWAS scan. It is important to note that our criteria to evaluate the estimation procedures differ from the hypothesis testing, and has a distinct focus on the estimation accuracy (e.g., measured by rooted mean square errors, or RMSE) on the gene-to-trait effects.

It might be intuitive to estimate the causal effect from a target gene to the complex trait of interest by simply regressing the complex trait phenotype on the composite IV or genetically predicted gene expression of the target gene. This approach, however, is not principled mainly because it ignores substantial uncertainty associated with the imputed gene expression levels. In practice, such *ad-hoc* approach shows both noticeable bias and large variance in estimation (Figure 3).

**Figure 3:**
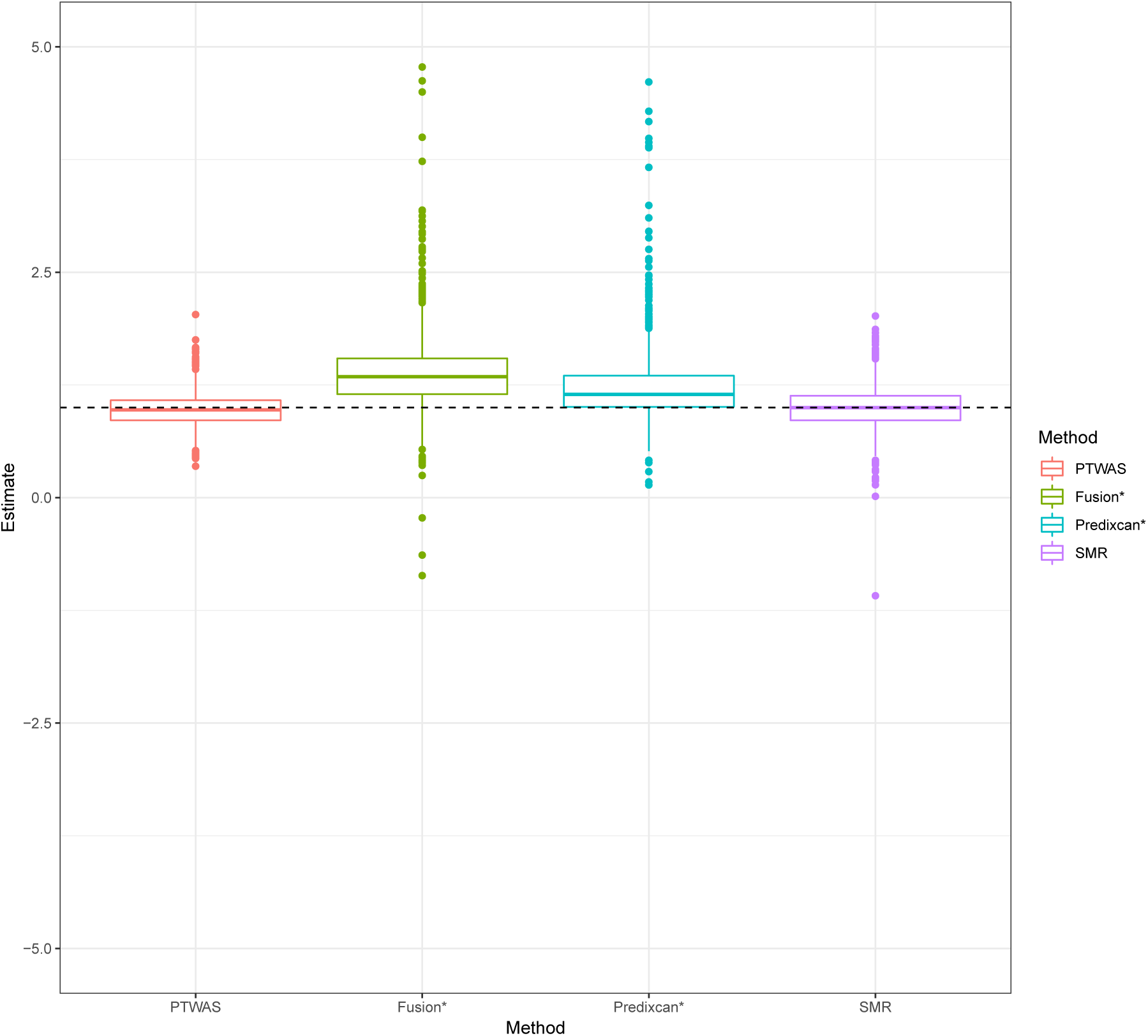
Boxplots of point estimates of causal effects by various methods in simulations. The true causal effect is set at 1.00 (dotted horizontal line) for all simulations. A total of 800 genes that pass the initial PTWAS scan (with *p*-value cutoff 0.05) are examined by each method. SMR utilizes the standard two-stage least squares algorithm based on the top eQTL SNP for each gene. For TWAS-Fusion and PrediXcan, the point estimates are obtained by regressing the phenotype data on the corresponding predicted gene expression levels. Note that effect size estimation is not the designed usage of TWAS-Fusion and PrediXcan. Our intention is to illustrate that methods designed for testing may not be suitable for estimation. Among all methods compared, PTWAS and SMR yield seemingly unbiased estimates. The results by PTWAS are, overall, more accurate.

We evaluate the proposed effect estimation procedure and compare its performance to SMR, which implements a standard two-stage least square (2SLS) algorithm by utilizing the strongest eQTL SNP (form single-SNP analysis) as the sole instrument. In our simulation studies, both PTWAS and SMR yield seemingly unbiased estimates (Figure 3). However, the results from PTWAS show higher accuracy. Even when a single (best) eQTL signal is utilized for estimation, we find that PTWAS is slightly more accurate (RMSE = 0.39) than SMR (RMSE = 0.44). This is likely due to the unique design of PTWAS that averages single-SNP estimates over a set of highly linked SNPs.

In all cases, we find that the estimation accuracy is directly correlated with the strength of instruments (i.e., eQTLs), which is quantified by the corresponding signal-level PIPs (Figure 4). By employing stronger eQTLs into the PTWAS estimation procedure, the accuracy monotonically increases. This observation directly verifies the estimation principle from the IV analysis. For a given SPIP threshold, we find that the estimates by combining multiple independent eQTLs (whenever available) tend to be more accurate than those relying on a single eQTL signal cluster. Furthermore, when the strength of the eQTLs is well controlled, we observe the causal effect estimates by independent eQTLs become highly consistent under the true causal model as illustrated by the average *I*^2^ in Figure 4. This particular observation validates our rationale for the model diagnosis procedure in PTWAS: the *I*^2^ computed from multiple independent eQTLs with modest strength (SPIP > 0.50) is expected to be less than 0.10.

**Figure 4:**
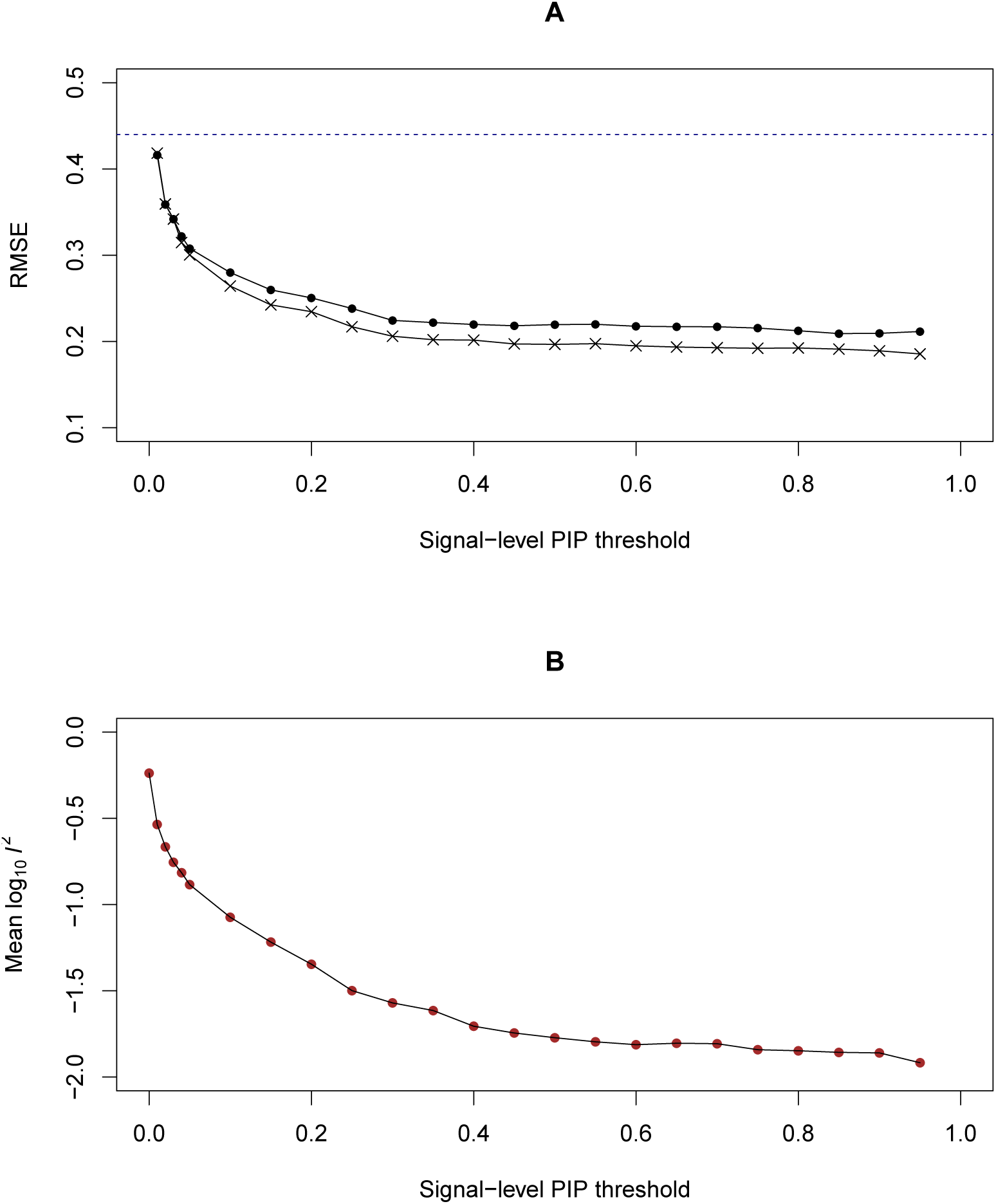
Impact of strength of eQTLs on causal effect estimation in PTWAS. 800 genes that pass the initial PTWAS scan (*p*-value ≤ 0.05) are used in this experiment. Panel A illustrates the relationship between the accuracy of the estimation (measured by rooted mean square error, RMSE) and the strength of the eQTL instruments (measured by signal-level PIP). The filled points represent all the genes satisfying the SPIP threshold. The stars represent the subset of genes where multiple independent qualifying eQTL instruments are available. The dotted line represents the RMSE by SMR. Overall, estimation accuracy increases when the strength of eQTL instruments improves and utilizing multiple instruments increases the accuracy of the estimation. Panel B shows the average heterogeneity of causal effects estimated from multiple

Last, we consider the possibility of conducting the scan procedure using the estimated causal effects as the SMR test. Specifically, we construct a *z*-statistic from the estimated causal effect and the corresponding standard error. We find this approach has slightly better power (0.725 at FDR 5% level) than the SMR test (0.713). However, both are markedly less powerful than the TWAS-Fusion (0.781), PrediXcan (0.784), and the proposed PTWAS scan procedure (0.800) at the same rejection threshold. We suspect this phenomenon is partially due to the difficulty in estimating the SNP-level standard errors (used by both SMR and the proposed estimation procedure). The estimated standard errors tend to be conservative in the lack of sufficiently large samples. Therefore, we do not recommend to replace the proposed PTWAS scan procedure with the SMR-like test.

In summary, we conclude that both principled effect estimation approaches, SMR and PTWAS, produce unbiased estimates in causal effect estimation. PTWAS shows improved precision by accounting for LD, the strength of the eQTL instruments, and the multiple independent eQTLs. The simulations also provide us some practical guidelines on the threshold values for strong instruments. Informed by the simulation results, we decide to admit eQTLs with signal-level PIP > 0.50 for model validation and effect size estimation in our real data analysis.

### 2.4 Multi-tissue PTWAS analysis of complex traits

We perform the PTWAS analysis on 114 complex traits from the UK Biobank (45 traits) and multiple large-scale consortia (69 traits), including CARDiOGRAM, PGC, and GIANT, by the GTEx GWAS working group [20]. A complete summary of the complex trait datasets are provided in Supplementary Table S1. We analyze the multi-tissue eQTL data from release 8 of the GTEx project [9] using DAP [11, 12] to generate the required probabilistic annotations of *cis*-eQTLs across 49 tissues. A total of 32,363 candidate genes are selected for the PTWAS analysis. For the analyses that we present in this paper, we utilize only the GWAS summary-level association statistics derived from the single-SNP analysis. Additional summary-level statistics for unreported GWAS SNPs in each trait are imputed to match the *cis*-SNPs in the GTEx panel.

#### 2.4.1 Scan trait-associated genes across multiple tissues

For each complex trait and each candidate gene, we first examine the global null hypothesis asserting that the target gene is not associated with the trait in any of the tissues. To this end, we construct the proposed PTWAS composite IVs for the target gene in all tissues and test their associations with the complex trait of interest. The resulting *p*-values from the single-tissue testing are subsequently combined using the ACAT method [21], which produces valid *p*-values for testing the global null hypotheses.

Across 114 traits, we reject the global null hypothesis for 115,327 gene-trait pairs at the false discovery rate (FDR) 5% level (at a more stringent *p*-value 0.05 level after Bonferroni correction, 30,777 gene-trait pairs are rejected). The complete list of significant gene-trait pairs is summarized in Supplementary Table S2 The genes rejected per trait have a notably skewed distribution (Figure 5). For well-known polygenic traits (e.g., height, BMI) and physiological traits (e.g., white blood cell count, lymphocyte count), a large number of genes are identified. The PTWAS analysis of the standing height data of UK Biobank identifies 13,696 genes at 5% FDR level, or 42% of total genes tested. On the other extreme, for 25 out of 114 traits, less than 10 genes are rejected. As expected, we find the power of discoveries is strongly correlated with the quality, sample size and power of the corresponding GWAS data as the composite IVs constructed from the GTEx data remain invariant for a given tissue across all traits. In the 45 traits from the UK biobank, we observe a strong correlation (ρ = 0.55, *p*-value = 1.1 × 10*^−^*^4^) between the TWAS discovery and the estimated heritability (data obtained from https://nealelab.github.io/UKBB_ldsc/). Figure 6 shows a comparison of multi-tissue PTWAS scan *p*-values of height-associated genes from UK biobank data and GIANT consortium. The strong linear trend indicates that both studies consistently give higher ranks of association for the same set of genes. However, the evidence from the UK Biobank data, with a much larger sample size, is often orders of magnitude stronger.

**Figure 5:**
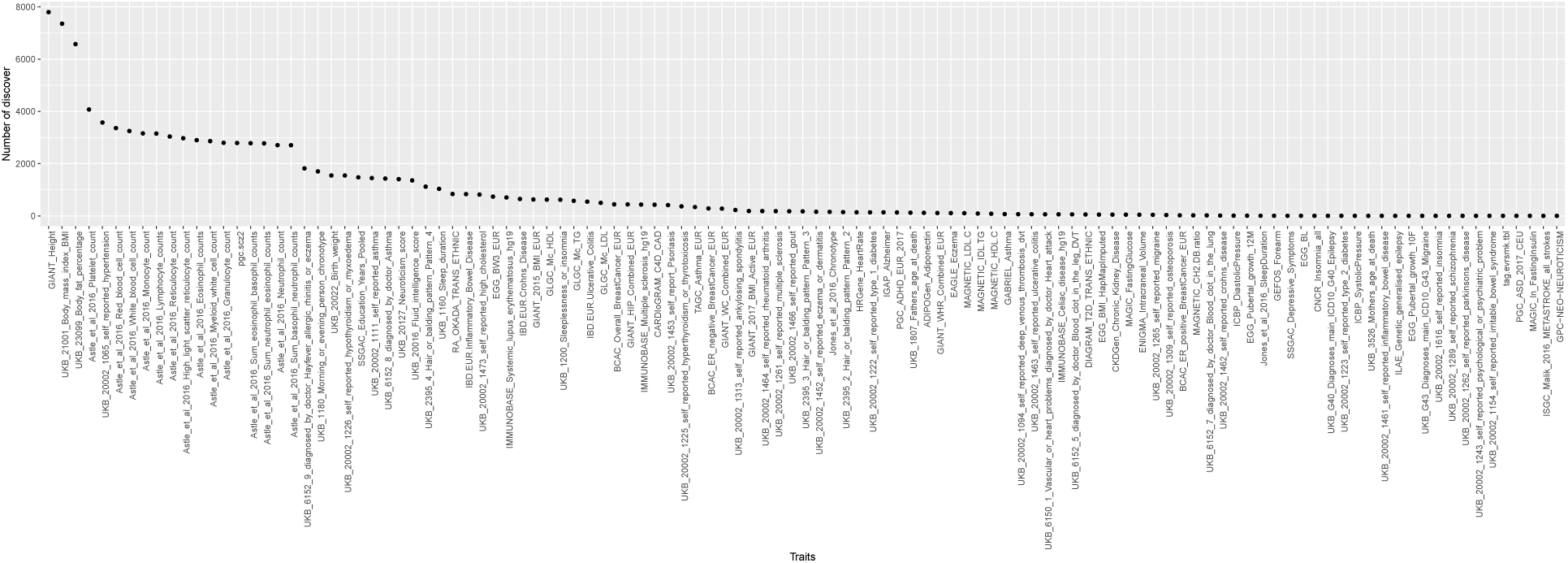
Discovery in PTWAS scan by traits. The results are from the analysis of GTEx eQTL data in 49 tissues and 114 complex traits using the PTWAS scan. Each point represents the number of rejected genes at the 5% FDR level for the corresponding trait. The distribution is highly skewed: a large proportion of tested genes are rejected in polygenic traits like heights, whereas few rejections are reported for > 20 traits.

**Figure 6:**
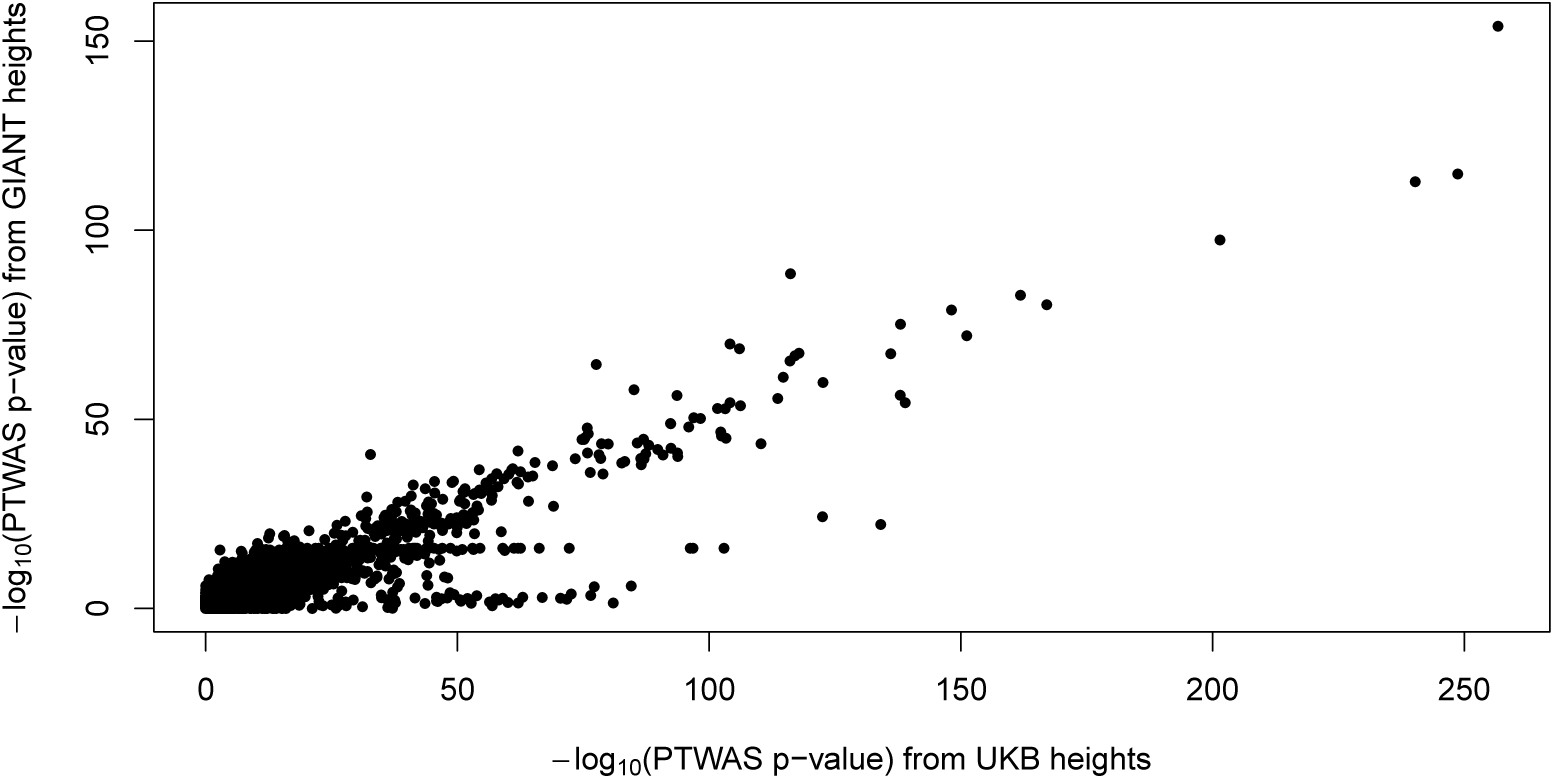
Comparison of PTWAS *p*-values for the height data from GIANT consortium and UK Biobank. Each data point represents a single gene whose PTWAS *p*-values are computed from GIANT and UK Biobank data, respectively. There is a strong linear trend for highly significant *p*-values (overall spearman’s correlation = 0.50, *p <* 2.2 × 10*^−^*^16^). However, UK Biobank data, which have a larger sample size, show consistently higher significance.

Additionally, we observe that PTWAS consistently identifies more candidate genes than the existing approaches, which re-confirms our finding in the simulation study. In the joint analysis of the height data from the GIANT consortium and the eQTL data from the GTEx whole blood, we find that PTWAS and S-PrediXcan identify 2,065 and 1,385 genes at 5% FDR level, respectively. (Both approaches utilize the algorithm implemented in S-PrediXcan to test the gene-level association, the difference in discovery is solely driven by the difference in computing the gene-level composite IVs/predicted gene expression levels).

#### 2.4.2 Validation of causal model assumption

We identify 24,548 unique genes from the 115,327 significant gene-trait pairs discovered at the cross-tissue scan stage. Within this set of genes, there are 401,889 gene-tissue pairs, corresponding to 2,094,658 trait-tissue-gene combinations, containing at least one cis-eQTL signals with SPIP ≥ 0.50, which we deem suitable for causal effect estimation. This set of gene-tissue pairs also show widespread allelic heterogeneity (Supplementary Figure S3). We further identify 124,858 gene-tissue pairs, corresponding to 664,868 trait-tissue-gene combinations that are suitable (i.e., each identified gene in the corresponding tissue has at least two independent eligible eQTLs) for the proposed causal model validation analysis. The complete results for all trait-tissue-gene combinations are summarized in Supplementary Table S3.

Within these testable instances, we observe that the majority show high levels of consistency among independent eQTL instruments (Figure 7). Specifically, we find 54.5% (or 362,315 trait-tissue-gene combinations) of the tested cases have *I*^2^ ≤ 0.05, while there are only 31.7% (or 211,291) of the cases with *I*^2^ values exceeding 0.5. As expected, we find that the ability to detect high degrees of heterogeneity in estimated causal effects improves with the increased availability of independent eQTL instruments. Nevertheless, the main feature of the *I*^2^ distribution remains the same for different numbers of available IVs (Figure 7).

**Figure 7:**
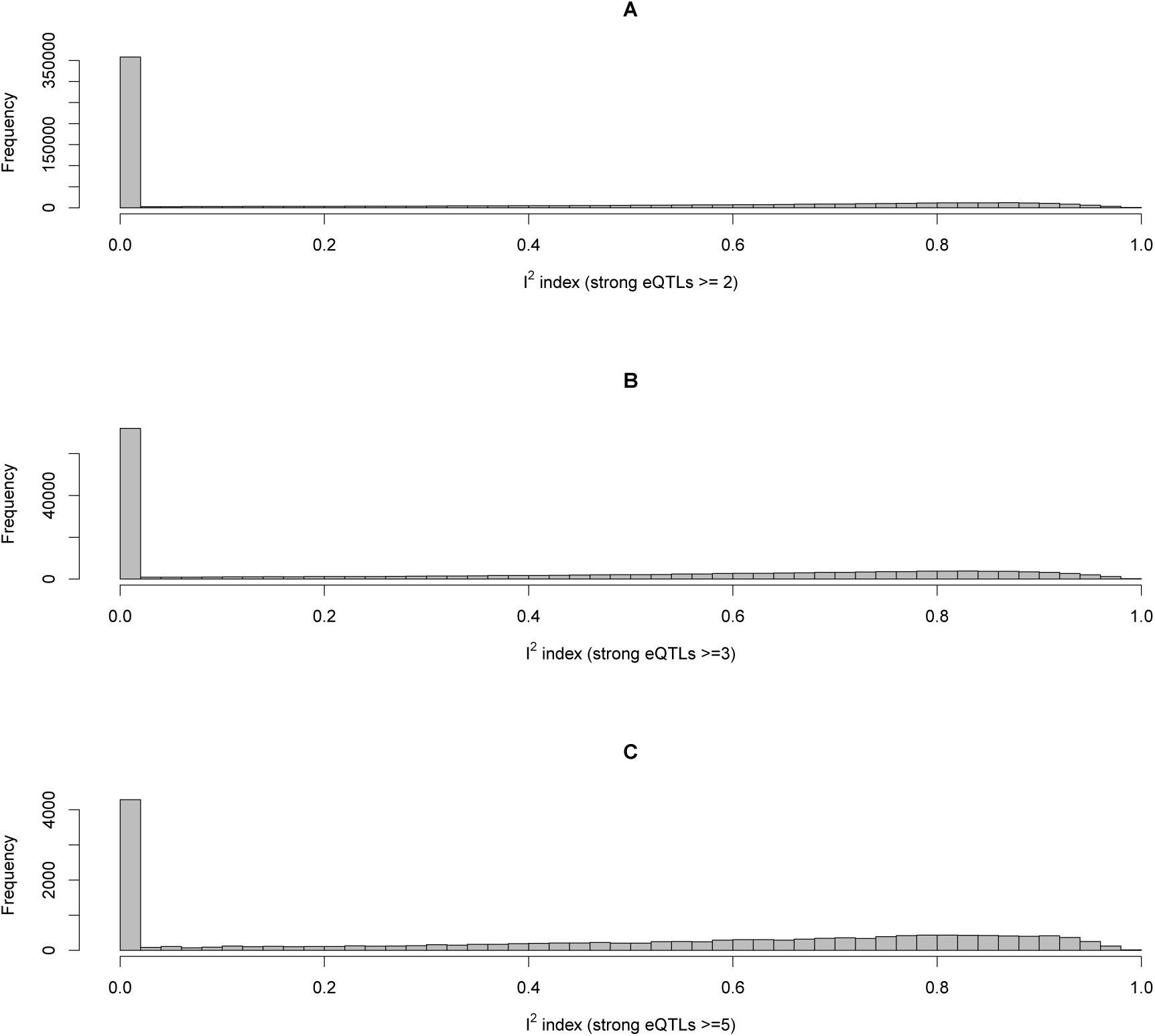
Histograms of *I*^2^ distribution for validating exclusion restriction from PTWAS scan signals. Panel A shows the histogram of the *I*^2^ statistics computed from 2.09 million trait-gene-tissue combinations, where the gene has at least two independent eQTLs with individual signal-level PIP > 0.50. For each trait-gene-tissue pair, the *I*^2^ index measures the consistency of the causal effects estimated from each independent eQTL. Panel B and C show the Histograms with at least 3 and 5 eligible independent eQTLs, respectively. The overall patterns of the distributions are the same in all panels, where majority of the test cases show strong consistency among independent instruments. However, the ability of identifying violations of ER improves with the increasing availability of independent eQTL instruments.

We show some examples of PTWAS scan signals that seemingly violate the exclusion restriction in Figure 8: *GUCY1A1* to cardiovascular disease (CAD) in artery tibial (*I*^2^ = 0.94), *CTB-171A8.1* to LDL cholesterol in artery aorta (*I*^2^ = 0.96), and *HLA-DQB2* to height in muscle skeletal (*I*^2^ = 0.98). (The specific tissues are where the most significant PTWAS scan *p*-values are identified.) In these examples, we observe the common pattern that the SNP-level estimates from correlated SNPs within each signal cluster are highly consistent, yet between the signal clusters, the estimates show high levels of inconsistency. In literature, *GUCY1A1* is considered a potential causal gene to CAD. Despite the association evidence discovered in a seemingly relevant tissue, the substantial inconsistency of the estimated gene-to-trait effects indicates that the role of the gene to the underlying disease can be complicated. In the case of the gene *HLA-DQB2* where we compute the *I*^2^ from the UK Biobank data, a similar high *I*^2^ value is also confirmed by the height data from the GIANT consortium (*I*^2^ = 0.77). Similarly in the case of *CTB-171A8.1* where the original *I*^2^ is computed from the LDL data from the global lipid genetics consortium (GLGC), a confirmative *I*^2^ = 0.97 is also found from the data of self-reported high cholesterol levels (binary outcomes) in the UK Biobank.

**Figure 8:**
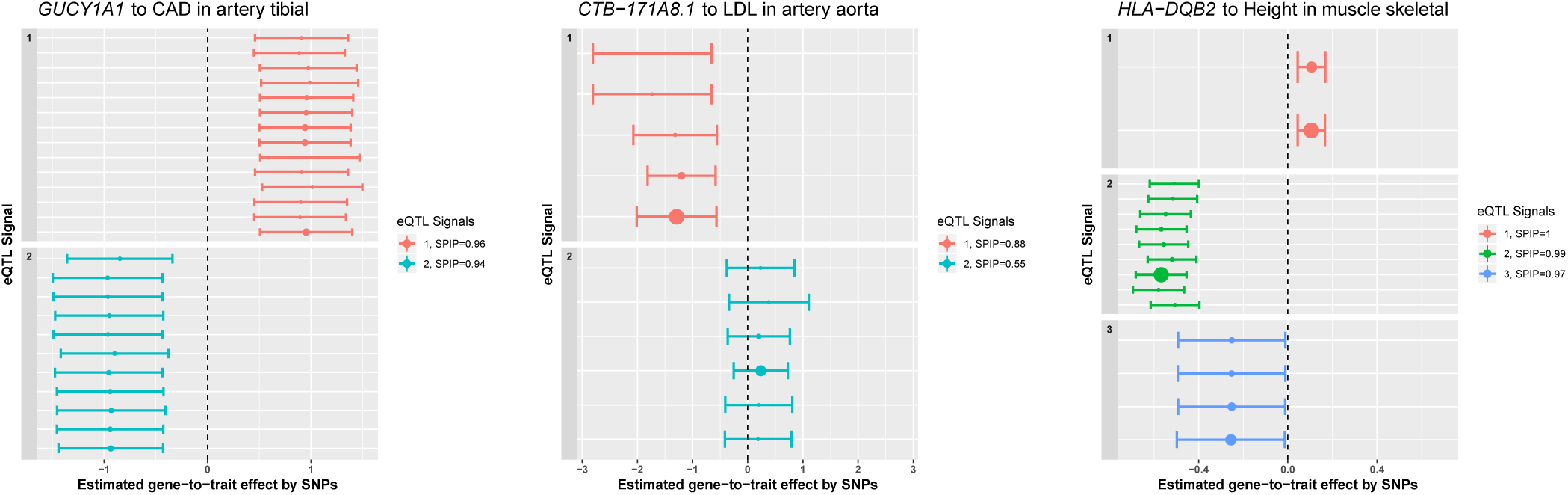
Examples of identified violations of exclusion restrictions. All three examples are identified from the PTWAS scan procedure, the corresponding tissues represent the tissues where the minimum *p*-values are achieved. Each SNP-level estimate and its corresponding 95% confidence interval are plotted. Data points with the same color represent the SNPs within the same signal cluster and in LD. The size of the point reflects the relative magnitude of SNP-level PIP. In all examples, within cluster estimates are highly consistent; estimates by different clusters are qualitatively different, which is a clear indication of violation of the exclusion restriction. The *I*^2^ statistics computed for the three cases are 0.94 (*GUCY1A1*), 0.96 (*CTB-171A8.1*), and 0.98 (*HLA-DQB2*), respectively.

#### 2.4.3 Gene effects to complex traits across multiple tissues

We proceed to estimate the potential causal effects from the genes implicated in the PTWAS scan to the complex traits of interest. For this analysis, we remove the 211,291 trait-gene-tissue combinations flagged in the ER validating analysis (*I*^2^ > 0.5) but include all the trait-gene-tissue combinations with only a single eligible eQTL instrument. In the end, we estimate the causal effects for 1,883,367 trait-gene-tissue combinations, corresponding to 114,083 gene-trait pairs, implicated by the PTWAS scan using the proposed estimation procedure. On average, each gene-trait pair is estimated in ~ 17 different tissues. The complete estimation results are summarized in Supplementary Table S3.

We examine the heterogeneity of the estimated gene-to-trait effects across multiple tissues, for which we again compute an *I*^2^ statistic (The complete results are summarized in Supplementary Table S4). The resulting *I*^2^ statistics display a clear bi-modal distribution (Figure S4). A significant proportion of the TWAS signals exhibit a high-level of consistency in effect sizes across different tissues: for all the investigated gene-trait pairs measured in multiple tissues, 46.9% show strong consistency with *I*^2^ < 0.1, whereas 12.8% exhibit a significant degree of heterogeneity with *I*^2^ > 0.75, suggesting potential gene-environment interactions (i.e., environmental factors modify the gene-to-trait effects). The median and the mean of all the cross-tissue *I*^2^ statistics are 0.30 and 0.18, respectively. Note that the tissue-specific effects of eQTLs drive the observed heterogeneity of the estimated gene-to-trait effects across tissues. The overall pattern that we observe from this analysis is qualitatively similoar to the reported tissue-specificity of *cis*-eQTLs in the literature: while a good proportion of eQTLs behaves in a tissue-consistent manner, a noticeable proportion shows strong evidence of tissue-specific eQTL effects [22, 23, 24]. This finding also confirms that, at cellular level, gene-environment interactions are widespread.

The estimated tissue-specific gene-to-trait effects provide potential insights for understanding the molecular mechanisms of complex diseases, many of which have been well-documented in the literature. In Figure 9, we highlight 4 examples: *SORT1* gene for LDL cholesterol, *CETP* gene for HDL cholesterol, and *PHACTR1, MRAS* genes for cardiovascular disease (CAD). In some cases, we find that the strongest or most confident effect size estimates are obtained from the biologically most relevant tissues, e.g., *SORT1* in liver and *PHACTR1* in artery coronary. In other cases, the estimated effects are similar in a group of similar tissues which possibly share some common cell types. The estimates also confirm the directional effects from the genes to corresponding traits: increased gene expression of *SORT1* and *PHACTR1* increase LDL levels and the risk of CAD, respectively; while increased gene expression of *CETP* and *MRAS* decrease the HDL level and the risk of CAD, respectively.

**Figure 9:**
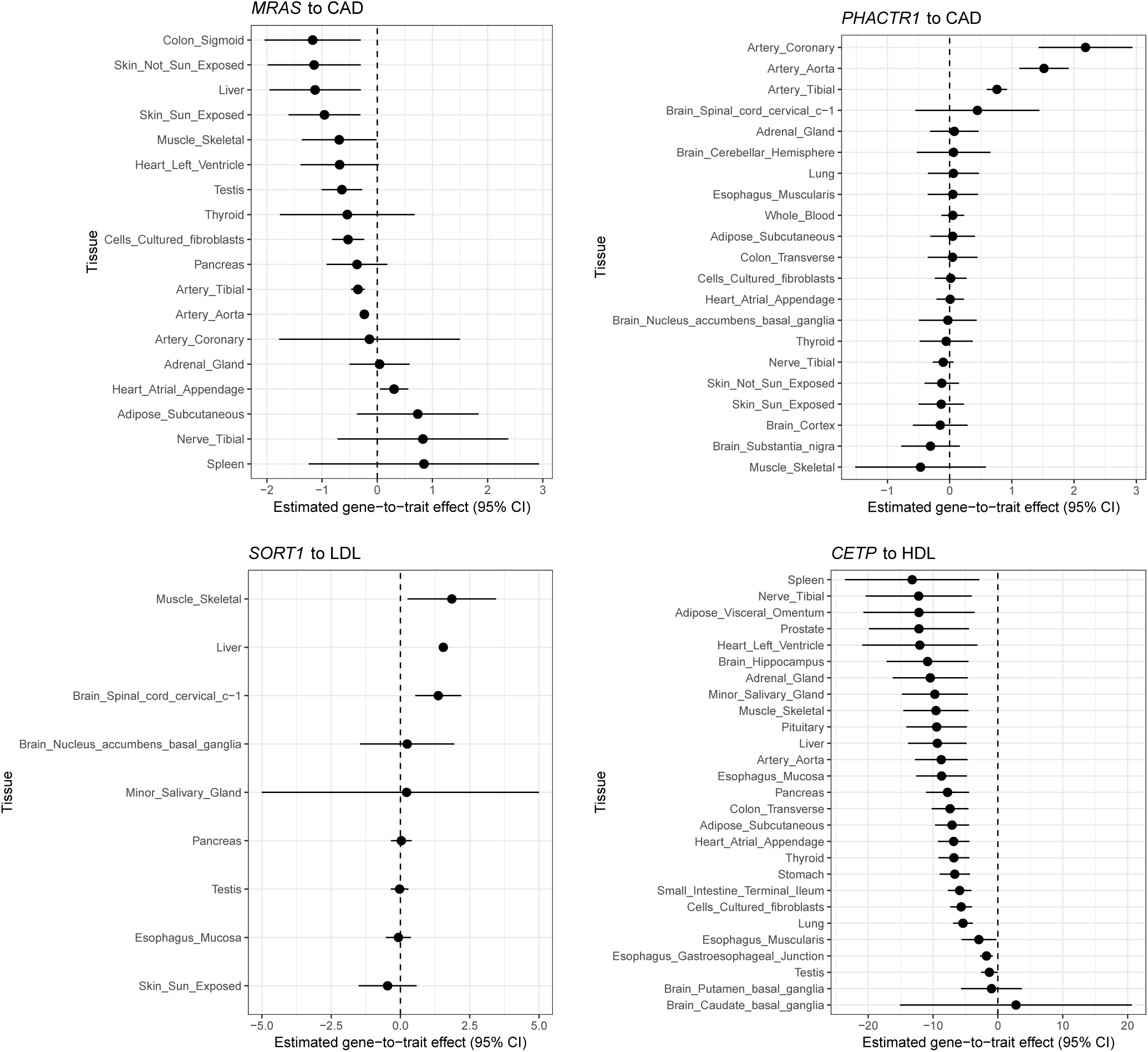
Examples of estimated gene-to-trait effects in different tissues. The selected gene-trait pairs have well-established causal relationships in literature. Here we observe different tissue-consistent and tissue-specific patterns in each gene-trait pair.

#### 2.4.4 Comparison to colocalization analysis

We compare the results from our multi-tissue PTWAS scan to the results from the colocalization analysis implemented in the software package ENLOC [25]. Colocalization analysis is a different type of integrative analysis approach that serves complementary goals as the TWAS analysis. Specifically, it aims to identify the overlapping of GWAS and eQTL signals at the single-variant level. Overall, the ENLOC analysis identifies much fewer gene-trait pairs where causal GWAS hits and *cis*-eQTLs are overlapping with high confidence. This is likely due to the fact that the GWAS and the GTEx data are from different cohorts (the two-sample design) and their LD patterns do not match exactly. The mismatch in LD patterns tends to increase the uncertainty of SNP-level colocalizations. Thus, relatively fewer signals can be assessed with very high colocalization probabilities.

Nevertheless, we observe that the gene-trait pairs identified by the PTWAS analysis are much more enriched with modest and high colocalization probabilities (Figure S5). The Kolmogorov-Smirnov statistic comparing the distributions of colocalization probabilities between all gene-trait pairs and PTWAS identified gene-trait pairs is 0.774 with the corresponding *p*-value < 2.2×10*^−^*^16^.

For a closer inspection, we examine the joint analyses of GWAS data of CAD from the CARDiOGRAM consortium and the GTEx eQTL data from whole blood. Figure 10 shows the scatter plot of the colocalization probabilities from ENLOC and the *p*-values from PTWAS. The plot displays a clear linear trend with high probabilities of colocalization corresponding to significant PTWAS *p*-values. However, the correlation is imperfect, as some most significant TWAS genes only exhibit weak evidence of colocalization, which is likely due to the differences of LD patterns in the GWAS and eQTL data for the relevant genomic regions.

**Figure 10:**
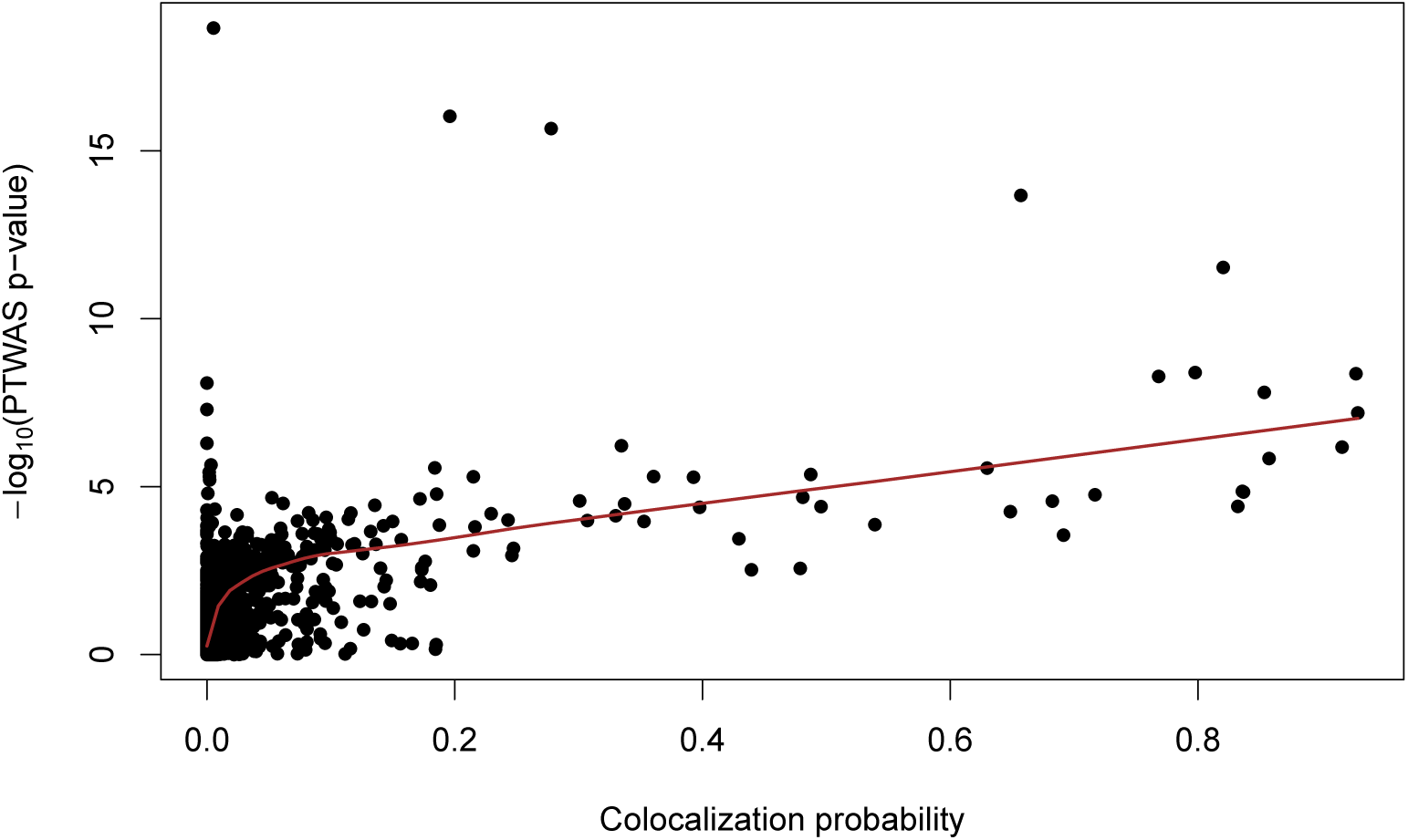
Comparison of −log10 *p*-values from PTWAS scan and gene-level colocalization probabilities in GTEx whole blood for the cardiovascular disease. Each data point corresponds a unique candidate gene. The red line is the fitted LOWESS curve, which indicates a strong positive linear correlation between the two quantities for significant PTWAS signals.

## 3 Discussion

In this paper, we propose a new computational framework to systematically investigate potential causal mechanisms from tens of thousands of candidate molecular phenotypes leading to complex diseases.

TWAS presents some unique challenges for the traditional causal inference methodologies like the IV analysis. Notably, the large number of candidate genes is rarely encountered in the application scenarios from econometrics and epidemiology. This unique feature prompts developing efficient multiple hypothesis testing procedures to screen candidate genes effectively. It is equally important to note that hypothesis testing may not be the endpoint of a sound causal inference: interpreting and validating the scan results have essential implications on follow-up studies. In this paper, we also illustrate that robust estimation of the causal effect, which has been emphasized in the traditional IV analysis, is critical to uncovering the impacts of cellular environments on disease etiology. Thus, we conclude that all three aspects of the PTWAS analysis: multiple hypothesis testing, model validation, and causal effect estimation, should be an integral part of causal inference in TWAS.

In PTWAS, our statistical strategies vary for different stages of the analysis. As noted by many authors [26, 16], although both hypothesis testing and causal effect estimation in the IV/MR analysis are operated based on the same set of casual assumptions, estimation procedures require additional parametric assumptions to formally define the casual effect (in PTWAS, we adopt the traditional definition of the causal effect via the linear structural equations embedded in the 2SLS method). Although it is possible to construct the test statistics based on the estimated gene-to-trait effects, we show that such test statistics tend to be underpowered compared to composite IV-based test statistics. (This is likely due to the overestimation of the corresponding standard errors.) The detailed discussion on this topic is outside the scope of this paper, and we will provide a full account in a follow-up technical article.

In this paper, we regard the commonly referred “genetic predictions of gene expressions” in the scan procedure as composite instrumental variables for investigating the causal relationships between candidate genes and the complex trait of interest. Although it is conceptually correct to interpret the composite IVs as genetic predictions of gene expression phenotype, we note that such predictions are generally inaccurate. The evidence here is two-fold: first, existing studies suggest the overall heritability of gene expressions, on average, is quite low [27]; second, as shown by many genomic experiments of differential gene expression analysis, manipulation of cellular environments can have drastic impacts on gene expression levels.

One of the advantages of the proposed PTWAS framework is its utilization of multiple independent eQTLs throughout the three stages of the analysis. This has been shown to improve the power of multiple hypothesis testing, enable the model validation, and increase the estimation efficiency. Importantly, our computational approaches carefully distinguish SNPs representing different independent eQTL signals and SNPs tagging the same eQTL signal. We note that regarding highly correlated SNPs as independent instruments are theoretically invalid. Instead, our approach uses a set of eQTL SNPs in LD (i.e., a Bayesian credible set) to represent an instrument and weight the evidence from individual SNPs through Bayesian model averaging (BMA). This strategy is proven effective in our simulations and real data analysis by carrying over the SNP-level uncertainty from the eQTL analysis. Recently, [28] propose an IV analysis approach, TWMR, to utilize multiple independent eQTLs identified from conditional analysis for estimating gene-to-trait effects in TWAS. Many authors have reported Bayesian multi-SNP fine-mapping methods, e.g., DAP, are more powerful than conditional analysis-based methods to effectively identify independent eQTLs [29, 30, 11]. Furthermore, TWMR estimation can be viewed as a special case of the proposed BMA weighting in PTWAS, which assigns all weights to the lead eQTL SNP within each signal cluster. However, such extreme weighting scheme seems sub-optimal in estimation accuracy as shown in our simulation studies.

Recently emerged TWAS methods, represented by FOCUS [31] and TWMR [28], also consider the strong correlations among the observed expressions of multiple genes. In the extreme cases, if the expressions of two genes are perfectly correlated, the true causal gene is not identifiable. Analyzing one gene at a time can fail to account for such correlations and result in over-reporting of causal genes. We have not explicitly addressed this issue in this paper due to our limited scope. However, we note the test statistics/*p*-values derived from our novel PTWAS scan procedure can be directly plugged into the fine-mapping methods implemented in FOCUS and yield desired credible sets of causal genes. We will continuously pursue this direction in our follow-up work.

Finally, we acknowledge that, like any observational data based causal inference approach, the proposed procedure is imperfect. It is mainly because some critical causal assumptions can not be rigorously verified, and some required conditions are not satisfied for every gene-trait pair in practice. In our application context, a good proportion of genes lack discoveries of multiple independent and strong eQTLs. Thus, the proposed model validation procedure is not applicable. The emergence of large-scale eQTL datasets will alleviate this problem in the near future as the ability to uncover multiple eQTLs is directly correlated with the sample size of eQTL [9]. Nevertheless, from a theoretical point of view, only severe violations of causal assumptions can be detected by model diagnosis approaches. In other words, passing the model validation procedure does not *prove* the assumptions are true but only indicate that they are reasonable given the data at hand. Should such caveat deter our efforts to apply the proposed approaches, or more generally, formulate and perform observational-data based causal inference in TWAS? Our opinion is no. The challenge is not unique to the field of genetics and genomics. We believe that our proposed approach falls into the category of “shoe leather” methodology advocated by David Freedman [32], which “*exploits natural variation to mitigate confounding and relies on intimate knowledge of the subject matter to develop meticulous research designs and eliminate rival explanations.*”

## 4 Methods

The probabilistic annotations of eQTLs used by PTWAS require comprehensive characterizations of the strengths and uncertainty of the genetic associations of expression traits at different levels. At the model level, each plausible association model (i.e., a combination of SNPs, denoted by *M_i_*) is assessed by a *posterior model probability*, *P_Mi_*. When multiple independent eQTLs coexist for a gene, the model including all relevant representing SNPs should have much higher posterior probability than the models including only single or incomplete eQTLs. A *SNP-level PIP*, *p_j_*, characterizes if a particular SNP *j* is a causal eQTL. In the presence of LD, causal eQTLs may not be statistically identifiable even if the evidence for the existence of an eQTL can be overwhelmingly strong [25]. Thus, we use the *signal-level PIP* to quantify the overall strength of an independent eQTL. Specifically for each potential eQTL signal *k*, we identify a set of SNPs in LD that represent the same association signal. The corresponding signal-level PIP is computed by *q_k_* =∑*_j∈Sk_ p_j_*. Thus, the comprehensive probabilistic annotations of eQTLs for a gene is given by ({*M_i_, P_M_i__*)}, {(*S_k_, q_k_*)}, {*p_j_*}. All these information are used for different tasks in PTWAS.

### 4.1 TWAS as IV analysis

The instrumental variable analysis can be represented by the graphical model in Supplementary Figure S1. There are three key assumptions to establish causal implications from the IV analysis. In the context of TWAS, they can be characterized as: i) *inclusion restriction:* the selected instrument (*G*), i.e., genetic variants, must be associated with the expression of the target gene (*X*); ii) *randomization assumption:* the instrument is (marginally) independent of confounders (*U*); and iii) *exclusion restriction:* the selected genetic variants affect the complex trait of interest (*Y*) only through the target gene. Notably, the inclusion restriction (IR) requires the instruments are eQTLs, and the exclusion restriction (ER) explicitly excludes the possibility of (horizontal) pleiotropy. Under these causal assumptions, the evidence of the association between *G* and *Y*, i.e., 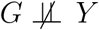, is sufficient to reject the null hypothesis that states no causal link between the target gene and the complex trait of interest [33, 26, 16].

The two-sample design refers to the setting that *G, X*, and *Y* are not taken from the same samples. Instead, the eQTL data (*G_x_, X*) and the GWAS data (*G_y_, Y*) are measured from two sets of non-overlapping samples. The two-sample design has some important implications on the IV analysis, especially the estimation bias of causal effects introduced by the weak instruments (i.e., weak eQTLs). In a one-sample design, the weak instruments can cause severe type I errors in testing causal relationships, whereas in a two-sample design, the bias is always towards 0, therefore, results only in the loss of power but not inflation of type I errors.

The weighted sum of IVs, i.e., ∑*_i_ w_i_G_i_*, is itself a valid instrumental variable because it satisfies all three IV assumptions. It is often referred to as a composite IV or allele score in literature [14, 15]. For hypothesis testing purposes, the aggregation of independent instruments with modest strength has an intuitive appeal for improved power for testing.

### 4.2 Composite IV for hypothesis testing

If the true causal eQTLs are known, the principled inference procedure in the IV analysis is the two-stage least squares (2SLS) method [16]. In the settings of the two-sample design, the first stage regression in the 2SLS finds the least-squares prediction of gene expressions using the eQTL data. The resulting prediction function is, by definition, a composite IV, where the weight for each individual genetic variant is the least square estimate of the corresponding genetic effect on the expression phenotype. In practice, even after the fine-mapping of eQTLs, there is substantial uncertainty regarding the number of independent eQTLs and the actual causal SNP for each eQTL. Selecting a single “best” association model from the eQTL data to perform the 2SLS procedure does not convey the uncertainty from the eQTL analysis, hence unlikely optimal.

We propose a *model averaging* approach that utilizes the posterior model probabilities {*P_Mi_*} to construct composite IVs based on the existing 2SLS procedure. For a set of *L* sparse candidate association models identified from the eQTL analysis, we fit each model *M_i_* by the least-square algorithm. We obtain an effect size estimate, 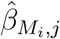, for each SNP *j* in model *M_i_*. (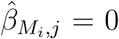xif SNP 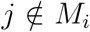) We then compute an overall weight for SNP *j* by averaging its estimated effects across all *L* candidate models, i.e.,

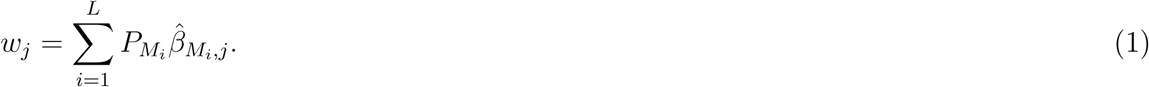

Finally, the proposed composite instrumental variable, *x*^, is computed by:

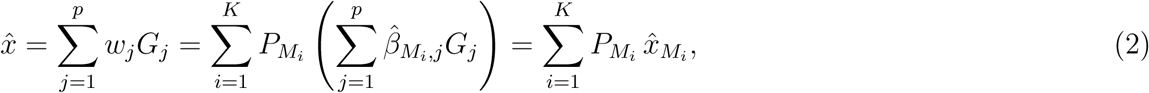

Where 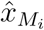 denotes the least squares prediction by the model *M_i_*. Therefore, the resulting composite IV can be naturally interpreted as an *ensemble prediction* of expression levels for the target gene (and each 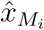 is known as a base prediction).

The composite IV (2) constructed by PTWAS has some unique and desired properties for the causal inference. First, it naturally considers multiple independent eQTL signals. Second, the weighting scheme also accounts for LD. Third, the proposed procedure provides a principled way to weigh weak versus strong instruments by utilizing the posterior model probabilities.

For a complex trait data set where individual-level data are available, we compute 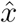 for each sample by plugging in the corresponding genotype data into the equation (2) and test its correlation with the observed complex trait phenotype. When only summary-level statistics (instead of individual-level genotype data) are made available from the complex trait data, we apply the approximate procedure described in [34], i.e., the S-PrediXcan algorithm, to compute the corresponding values of 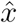 based on *z*-statistics.

### 4.3 Estimating causal effects

We take a slightly different strategy to estimate the causal effects from a target gene to the trait of interest to ensure accuracy and robustness. Notably, we emphasize in obtaining effect size estimates from strong instruments, where we quantify the strength of an instrument by its corresponding signal-level PIP. The designed filtering of weak instruments is to avoid biased estimates. When multiple independent eQTLs are available, we aggregate the estimates from the individual eQTLs using a fixed-effect meta-analysis procedure.

The proposed estimating procedure starts with selecting strong eQTLs by thresholding on the corresponding signal-level PIPs. For a qualified eQTL signal with *m* member SNPs and signal probability 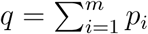, we then re-normalize the SNP probabilities by

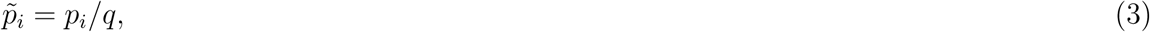

which corresponds to the conditional probability of SNP *i* being a causal eQTL. Next, we obtain an estimate of the causal effect, 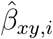 and the corresponding variance 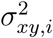 based on the summary statistics of SNP *i* using the standard 2SLS method (i.e., similar to SMR). Subsequently, by the law of total expectation, we combine the individual estimate from each member SNP weighted by its conditional causal probability, i.e.,

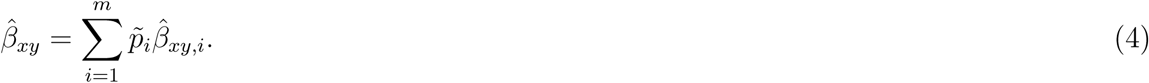

Similarly, by the law of total variance, we obtain

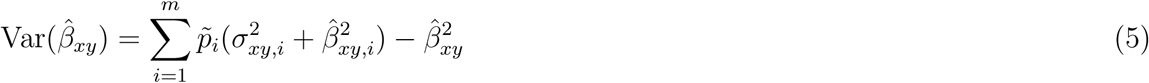

For each independent instrument/eQTL, the proposed estimator provides an intuitive and principled approach to fully account for LD.

Under the IV assumptions, each independent eQTL provides an independent estimate of the underlying causal effects. If the causal link from the target gene to the complex trait is indeed true, the multiple estimates from independent eQTLs are analogous to the multiple estimates from a meta-analysis, where the underlying true causal effect should be constant regardless of the instruments used for measurement. By this logic, it becomes natural to adopt a fixed-effect meta-analysis model to combine the individual 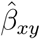 using the inverse variance weighting (IVW) scheme.

### 4.4 Diagnosis of exclusion restriction

The causality implications inferred from TWAS analysis are subject to the validity of the IV assumptions, especially the exclusion restriction (ER). Although rigorous validation of ER based on observed data is considered theoretically implausible [35, 26, 16], it is practically feasible to detect *severe departures* from ER when multiple independent instruments become available. If ER holds, the causal effects estimated from multiple independent eQTLs (using the procedure in the above section) should be highly consistent, and low levels of heterogeneity among estimates are expected. In contrast, observations of elevated heterogeneity levels should flag potential violations of ER.

To implement this idea, we compute an *I*^2^ statistic [17] based on the causal effect estimates from multiple independent eQTLs that are available. Namely,

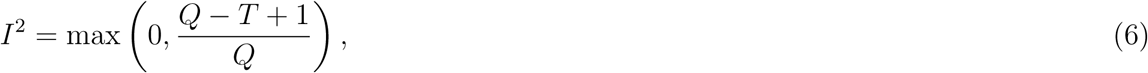

where *T* represents the number of independent eQTLs used for effect estimation, and *Q* is the Cochrans Q statistic and is given by

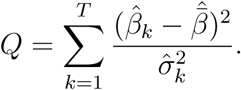

Specially, 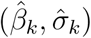 are estimated effect size and the corresponding standard error from the *k*-th independent eQTL, and 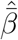 is the fixed-effect estimate of the overall causal effect. The *I*^2^ statistic ranges from 0 to 1 and is designed initially to represent the percentage of the variance observed in a meta-analysis that can be attributed to the heterogeneity among participating studies. In the application context of TWAS, *I*^2^ → 0 indicates reasonable consistency among estimates by multiple eQTLs, whereas *I*^2^ → 1 implies severe departures from ER.

We note that the underlying idea of the proposed model diagnosis approach is similar to the intuition behind Egger regression [8] that is widely applied in the context of MR analysis of multiple complex traits. However, the available independent eQTLs/instruments for any given gene are typically limited, which render the regression-type of diagnosis like Egger regression implausible. The proposed approach directly addresses this difficulty and is tailored for the application in TWAS.

We use *I*^2^ as a quantitative metric to measure the effect size heterogeneity in a comprehensible scale. It is theoretically possible to set up a hypothesis testing procedure to select genes with consistent eQTL-level estimates. Note that in this case, the null hypothesis has to assert heterogeneity in the underlying effect sizes. We have not seen any successful applications of such hypothesis testing. In practice, some authors suggest setting up the null hypothesis as in the Cochran’s *Q* test (which states all underlying effects are the same) and attempting to “*accept*” the null hypothesis. This, in our view, is a statistical mistake.

### 4.5 Simulation details

Throughout our simulation studies, we use the real genotype data from 706 whole blood samples of the GTEx v8 data to simulate expression and complex trait phenotype data. To mimic the two-sample design, we randomly select 400 individuals and 1000 genes to simulate their expression data. The remaining 306 individuals are used to simulate complex trait data.

For each gene, we select 1,500 *cis*-SNPs and independently sample the causal eQTLs from a Bernoulli distribution with the frequency = 0.002, Thus, on average, each gene has three causal eQTLs. The effect size of each causal gene is sampled from a Gaussian distribution with mean 0 and variance 0.5. We also add a polygenic background effect to all candidate SNPs, such that the heritability of the expression phenotype is roughly 50% (variations are due to the allele frequencies of the causal SNPs). More specifically, for individual *i*, the expression level for a candidate gene, *x_i_*, is simulated by

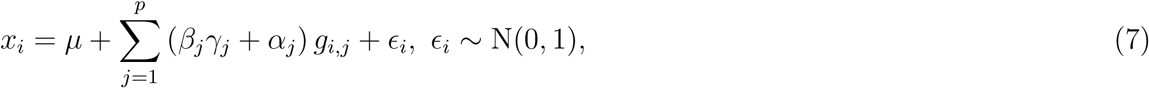

where *g_i,j_* represents the genotype of SNP *j* for the *i*-th individual, *β_j_*, *α_j_* represents the (strong) sparse genetic effect and the polygenic effect, respectively, and *γ_j_* is the latent Bernoulli random variable. The corresponding complex trait for individual *i*, *y_i_*, is subsequently simulated by a simple structural equation,

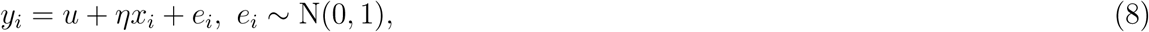

where *η* represents the causal gene-to-trait effects. To investigate the power and type I error control of the PTWAS scan procedure, we sample *η* from a Gaussian distribution *N* (0, 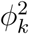) where the variance parameter is chosen from the set {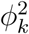 : 0, 0.2, 0.5, 0.8, 1, 1.2}. (Note that 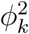 = 0 represents the null model). To examine the effect size estimation, we fix *η* = 1 for all simulated data sets. We simulate the expression data for all 706 individuals but use only the subset of 400 individuals for eQTL analysis. We only simulate the complex trait data for the subset of 306 individuals.

For the proposed PTWAS procedure, we analyze the eQTL data and construct the composite IVs using the software package DAP. For PrediXcan method, we apply the elastic net algorithm with cross-validation to find the genetic predictions of gene expressions. We also run the software package BSLMM to produce the predicted gene expression functions for TWAS-Fusion. When running the BSLMM, we apply the default settings except resetting the number of sampling steps to 200,000 and the burn-in steps to 100,000 for computational considerations.

### 4.6 Multi-tissue TWAS analysis

For multi-tissue TWAS analysis, we utilize the GTEx *cis*-eQTL data from 49 tissues (version 8) and 114 complex traits from the UK Biobank and other consortia. We perform multi-SNP fine-mapping of *cis*-eQTL analysis for each GTEx tissue using DAP, and generate the all three required levels of probabilistic annotations (i.e., posterior model probabilities, SNP-level PIPs, and signal-level PIPs) for the proposed PTWAS analysis.

#### 4.6.1 Data preparation

The 114 complex trait data sets are selected and harmonized by the GTEx GWAS analysis sub-group. For each complex trait, only summary statistics are extracted, and additional summary statistics are imputed for the GTEx SNPs that are not directly available from the original studies. The details of the GWAS data processing are documented in [20].

The probabilistic eQTL annotations required by PTWAS are derived from the GTEx release v8 data across 49 tissues using DAP [11, 12]. The pre-processing of the RNA-seq and genotype data follow the protocols of the GTEx data processing, which are detailed in [9]. The multi-SNP fine-mapping analysis by DAP uses the individual-level genotype data and controls the same set of covariates and PEER factors as in GTEx v8 single-SNP eQTL mapping.

#### 4.6.2 Multi-tissue PTWAS scan

Let 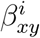 denotes the unobserved causal effect from gene *x* to trait *y* in the *i*-th tissue. We apply a multi-tissue PTWAS scan to test a global null hypothesis that states

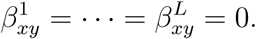

To carry out the testing procedure, we first derive a *p*-value, 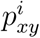, by applying the PTWAS scan procedure in the *i*-th tissue for the target gene-trait pair. We then apply the ACAT method [21] to the set of *p*-values for each gene-trait pair, {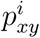 : 1, …, 49}, and compute a *p*-value for testing the global null hypothesis. For each trait, we finally apply Storey’s *q*-value procedure to control the false discovery rate at 5% level and identify the genes that reject the global null hypothesis.

#### 4.6.3 Estimating gene-to-trait effects

For each complex trait, we focus on the set of genes implicated from the PTWAS scan procedure. For a gene rejected at the scan stage (at FDR 5% level) for the trait of interest, we consider all its gene-tissue pairs (not just the most significant tissue) for the effect estimation and the model validation. We then select the gene-tissue-trait combination that contain at least one eQTL signal cluster with SPIP ≥ 0.50 for estimation purpose. This particular threshold is mainly informed by our simulation studies, where we observe reliable estimation results for signals with SPIP ≥ 0.50. To validate ER assumption, we require the selected gene-tissue-trait combinations to have more than two eQTL signal clusters with SPIP ≥ 0.50.

For each selected gene-tissue-trait combination, we carry out the proposed effect estimation and/or model validation procedures separately.

### 4.7 Software and data resources

The Github repository https://github.com/xqwen/ptwas/ contains the implementations of the algorithms described in this paper. The repository also provides a step-by-step guidelines on running the complete PTWAS analysis. The probabilistic annotations of eQTLs are generated by the software package DAP (https://github.com/xqwen/dap/) with both C++ and R implementations.

We have made the following resources available.

1. Probabilistic annotation of GTEx v8 *cis*-eQTLs across 49 tissues: https://storage.googleapis.com/gtex_analysis_v8/single_tissue_qtl_data/GTEx_v8_finemapping_DAPG.tar
2. Weights computed from GTEx v8 data for PTWAS scan across 49 tissues: https://tinyurl.com/yxe9k6vl
3. Summary statistics of 114 complex traits in GAMBIT format: https://tinyurl.com/y2hfhehj

## Supplementary Figures

**Figure S1:**
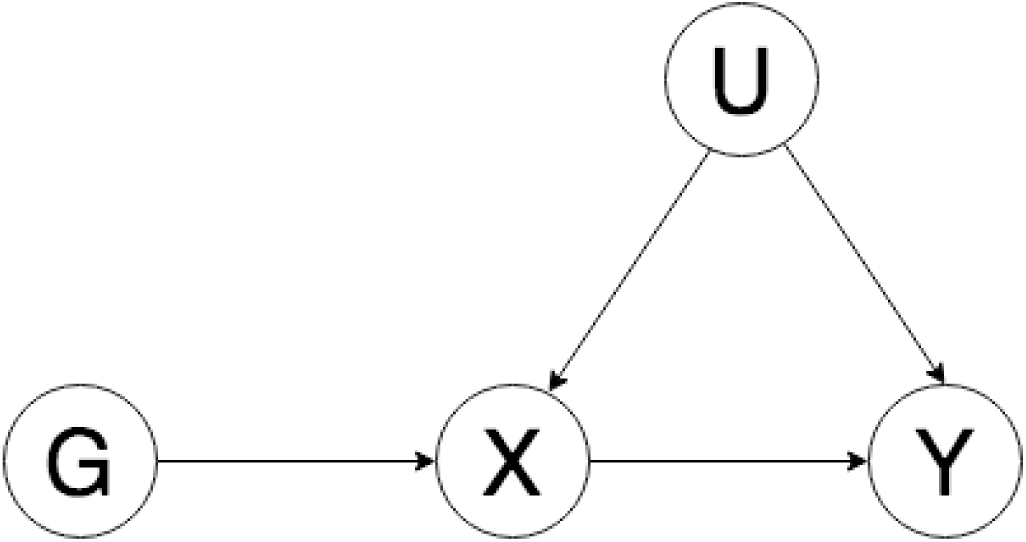
Diagram representing instrumental variable analysis. Variables *G, X, Y,* and *U* represent eQTLs, gene expressions, complex traits and unobserved confounding factors, respectively. The arrow from *X* to *Y* represents the causal relationship of interest.

**Figure S2:**
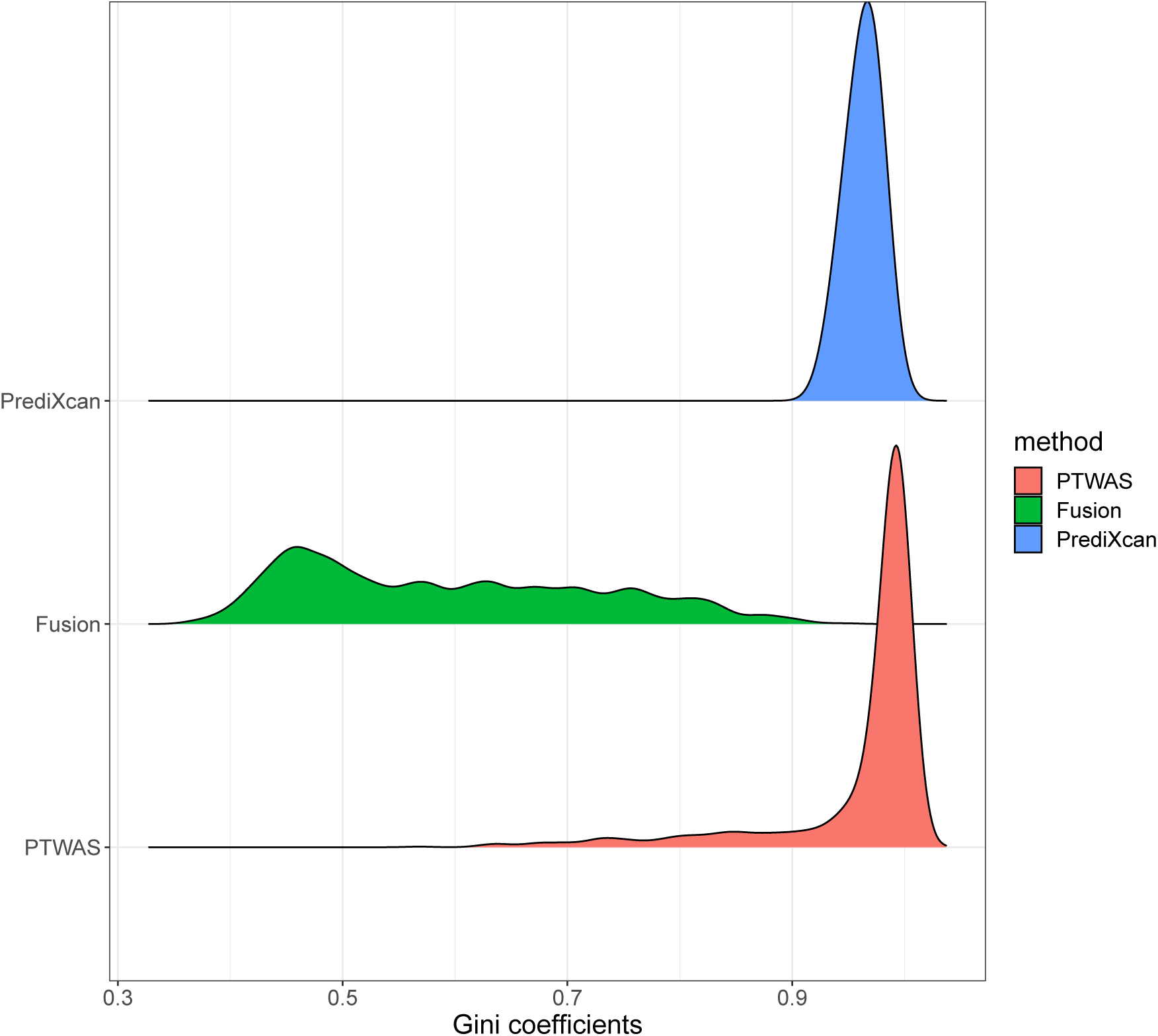
Distribution of Gini coefficients summarizing the sparsity of the composite IVs in different TWAS scan methods. For each simulated gene, we compute the Gini coefficient based on the assigned weights of all the *cis*-SNPs for computing the composite IV/predicted gene expression levels. A low (i.e., → 0) Gini coefficient indicates many SNPs play important roles; whereas a high value (i.e., → 1) suggests only a few SNPs make contributions. Among the methods compared, PrediXcan (based on ElasticNet algorithm) utilizes the most sparse set of SNPs for predicting gene expressions, and TWAS-Fusion utilizes many more SNPs (most of which are likely weak IVs). PTWAS is overall similar to PrediXcan but with a notable long left tail. This is mainly because the weight assignment in PTWAS takes accounts of LD: strong eQTL SNPs that are highly correlated (hence not identifiable) are assigned to comparable weights.

**Figure S3:**
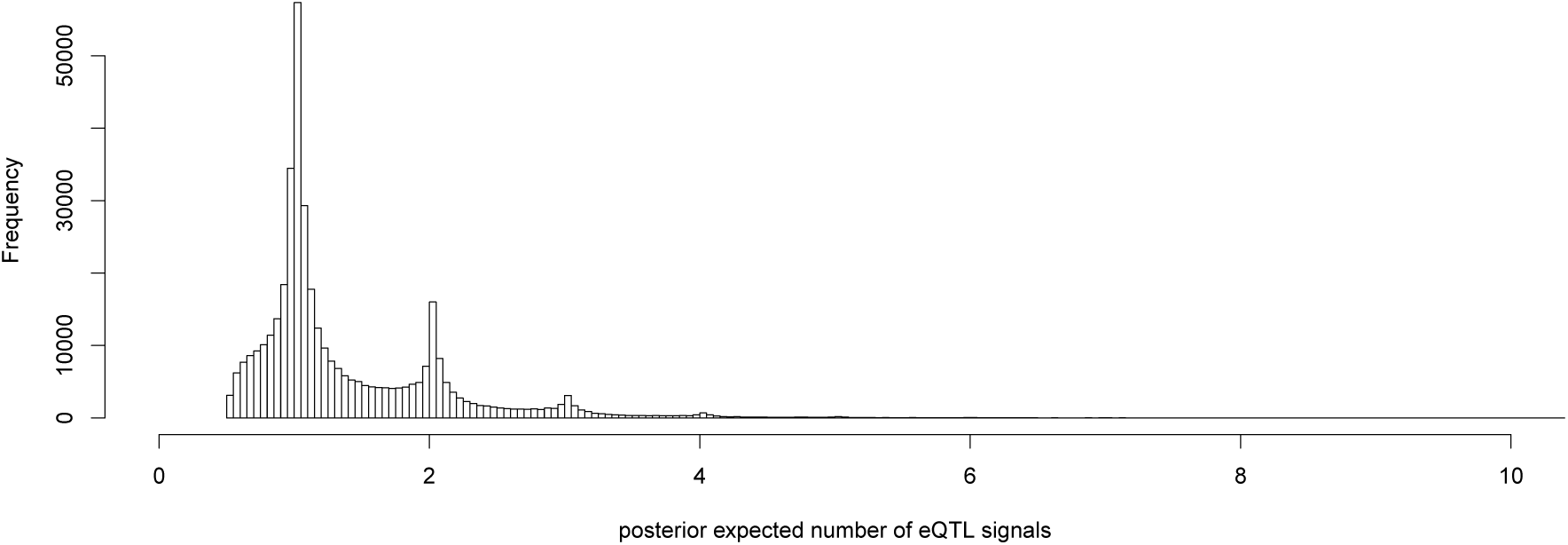
Allelic heterogeneity in implicated PTWAS signal genes. The histogram shows the distribution of the posterior expected number of *cis*-eQTLs of each unique gene-tissue pair that are implicated in the PTWAS scan and suitable for effect size estimation from the analysis of GTEx data and 114 complex traits. The plot indicates that a substantial proportion of gene-tissue pairs have more than 1 strong eQTLs.

**Figure S4:**
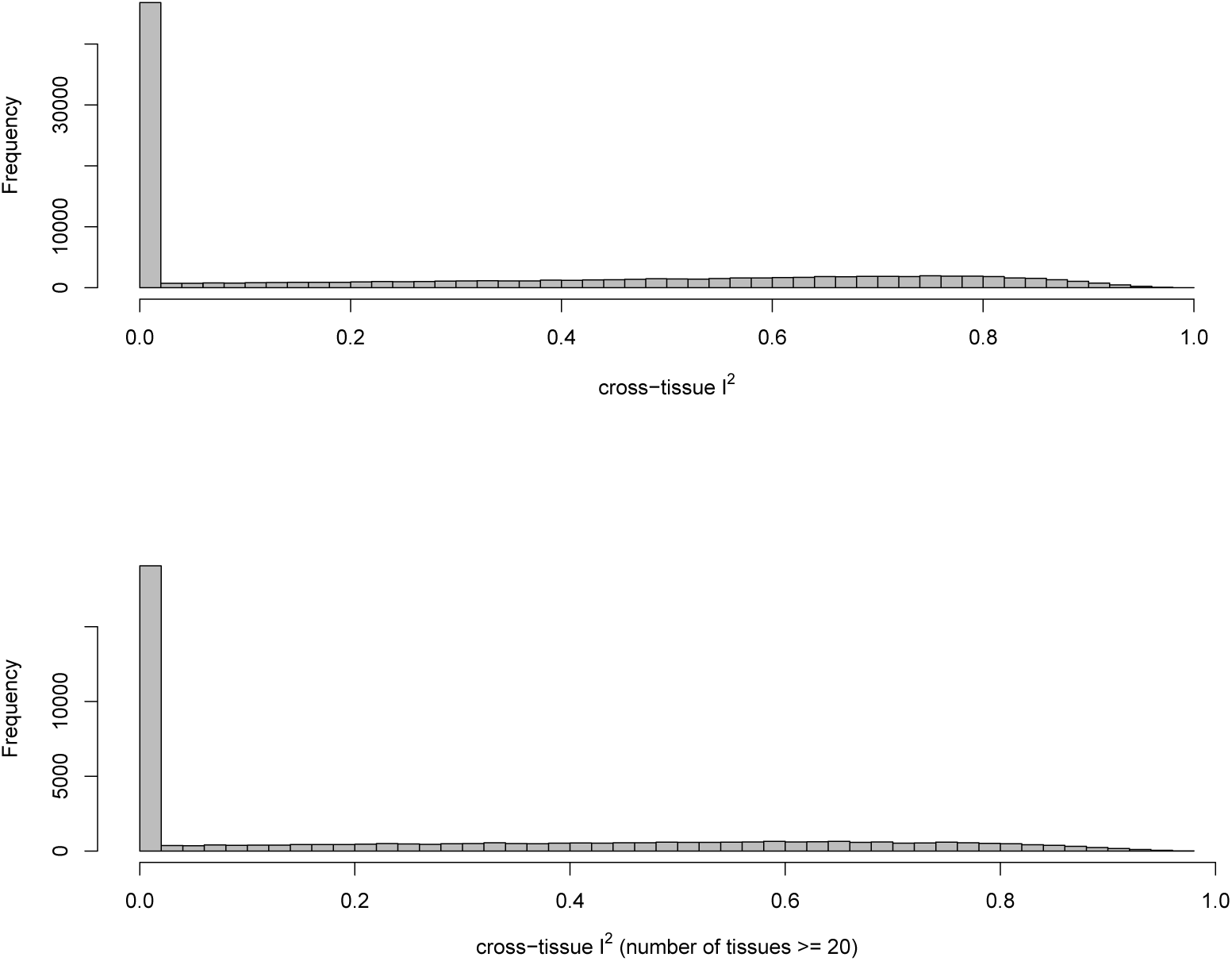
Heterogeneity of gene-to-trait effects across tissues. The histograms show the distributions of *I*^2^ statistics which quantify the heterogeneity of estimated gene-to-trait effects across different GTEx tissues. The top panel shows all eligible gene-trait paris. The bottom panel shows the gene-trait pairs that are measured in ≥ 20 different GTEx tissues. The overall patterns remain the same.

**Figure S5:**
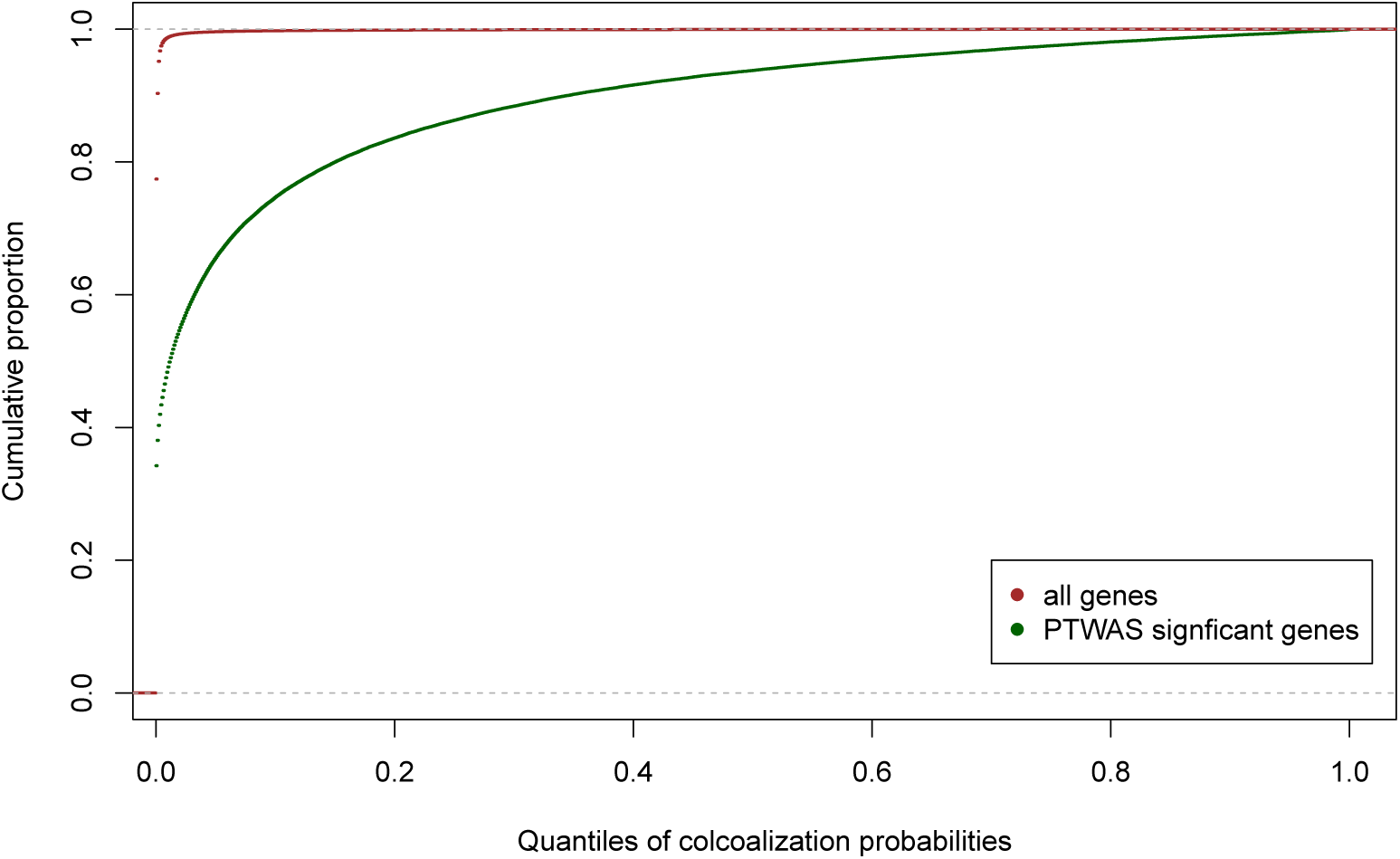
Comparison of distributions of colocalization probabilities between all genes and PTWAS significant genes. The red line represents the cumulative distribution function (cdf) of colocalization probabilities summarized from all 32,363 candidate genes in all 49 tissues across 114 traits. The green line represents the cdf from the corresponding PTWAS significant genes. It is clear that PTWAS signal genes are enriched with modest to high colocalization probabilities.

## Supplementary Tables

[See attached excel file]

**Table S1:** Descriptions of 114 complex trait datasets used in the PTWAS analysis.

[See attached text file]

**Table S2:** Complete list of significant gene-trait pairs at 5% FDR level identified in the multi-tissue PTWAS scan of 114 complex traits.

[See attached compressed text file]

**Table S3:** Complete list of estimation and model validation results for all gene-trait-tissue combinations.

[See attached text file]

**Table S4:** Complete cross-tissue heterogeneity results for all gene-trait pairs.

